# Dual PAX3/7 transcriptional activities spatially encode spinal cell fates through distinct gene networks

**DOI:** 10.1101/2025.03.11.642668

**Authors:** Robin Rondon, Théaud Hezez, Julien Richard Albert, Bernadette Drayton-Libotte, Gloria Gonzalez-Curto, Frédéric Relaix, Claire Dugast-Darzacq, Pascale Gilardi-Hebenstreit, Vanessa Ribes

**Affiliations:** Université Paris Cité, CNRS, Institut Jacques Monod, F-75013 Paris, France; Univ Paris Est Créteil, INSERM, EnVA, EFS, IMRB, F-94010 Créteil, France

**Keywords:** Neural fate specification, Transcriptional duality, Pioneer factors, PAX3/PAX7 regulatory networks, Spinal cord patterning, Chromatin dynamics, Enhancer-silencer interplay, BMP signalling, Functional genomics in organoids, CUT&Tag epigenomic profiling

## Abstract

Understanding how transcription factors regulate organized cellular diversity in developing tissues remains a major challenge due to their pleiotropic functions. We addressed this by monitoring and genetically modulating the activity of PAX3 and PAX7 during the specification of neural progenitor pools in the embryonic spinal cord. Using mouse models, we show that the balance between the transcriptional activating and repressing functions of these factors is modulated along the dorsoventral axis and is instructive to the patterning of spinal progenitor pools. By combining loss-of-function experiments with functional genomics in spinal organoids, we demonstrate that PAX-mediated repression and activation rely on distinct *cis-*regulatory genomic modules. This enables both the coexistence of their dual activity in dorsal cell progenitors and the specific control of two major differentiation programs. PAX promotes H3K27me3 deposition at silencers to repress ventral identities, while at enhancers, they act as pioneer factors, opening and activating *cis-*regulatory modules to specify dorsal-most identities. Finally, we show that this pioneer activity is restricted to cells exposed to BMP morphogens, ensuring spatial specificity. These findings reveal how PAX proteins, modulated by morphogen gradients, orchestrate neuronal diversity in the spinal cord, providing a robust framework for neural subtype specification.

## Introduction

The generation of an organized array of functionally distinct cell types is a fundamental outcome of development, with profound implications for the physiology of adult organs (Ramos et al. 2024; Lai et al. 2016). Transcription factors (TFs), through their ability to activate or repress gene expression, play a pivotal role in patterning cell fates within embryonic tissues (Spitz and Furlong 2012; Sagner and Briscoe 2019). However, for many TFs, the mechanisms by which their pleiotropic activity at *cis-*regulatory modules (CRMs) drives position-specific differentiation within a tissue remain poorly understood (Spitz and Furlong 2012; Sagner and Briscoe 2019). This challenge is further compounded by the fact that key TFs exhibit broad expression across cells destined to commit to multiple distinct lineages (Spitz and Furlong 2012; Koromila and Stathopoulos 2017).

The developing mammalian spinal cord serves as an ideal model to investigate TF pleiotropic activity due to its remarkable cellular diversity and the significant progress in understanding the role of TFs in regulating this diversity (Sagner and Briscoe 2019). In this tissue, two opposing morphogen gradients—Sonic Hedgehog (Shh) emanating from the ventral side and bone morphogenetic proteins (BMPs) and Wnts from the dorsal side— provide positional cues that guide the spatially organized expression of TFs in distinct, well-defined bands along the dorsoventral (DV) axis (Figure 1A)(Sagner and Briscoe 2019). The combined activity of these TFs defines eleven pools of neurogenic progenitors organized along this axis, each of which is committed to differentiation programmes into specific neuronal subtypes (Figure 1A)(Sagner and Briscoe 2019). This transcriptional regulation ensures the proper formation of locomotor and somatosensory circuits in the adult spinal cord (Sagner and Briscoe 2019). Among these eleven progenitor pools, six are located in the dorsal part of the spinal cord, designated as dp1 to dp6 in order from the most dorsal to the most ventral, and give rise to dI1 to dI6 interneurons (INs)(Figure 1A)(Sagner and Briscoe 2019). The remaining five pools are found in the ventral spinal cord, labelled p0, p1, p2, pMN, and p3, and generate V0 to V3 INs and motoneurons (MNs) (Figure 1A)(Sagner and Briscoe 2019).

**Figure 1.**
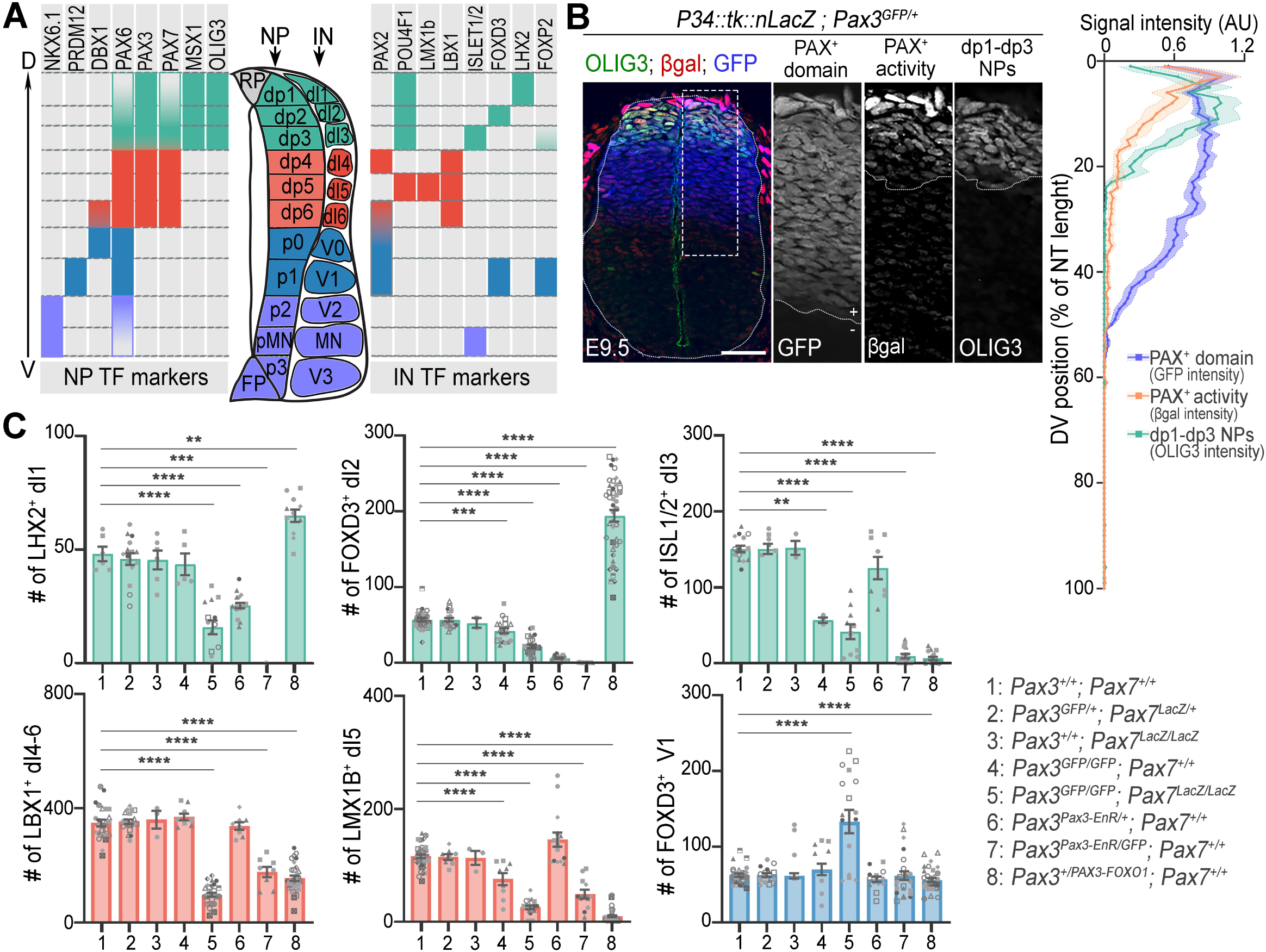
Neuronal diversity within the spinal cord of mouse embryos following the modulation of PAX3 and/or PAX7 transcriptional activity. (A) DV expression patterns of neural progenitor (NP) and interneuron (IN) TF markers. (B) Images: Immunostaining for β-galactosidase (β-gal, red/grey), OLIG3 (green/grey) and GFP (blue/grey) on a transverse section of E9.5 *P34::tk::LacZ; Pax3^GFP/+^* spinal cord at brachial level. Right panels are blown up of the squared region shown in the left panel. Graph: Quantification of the βgal, OLIG3 and GFP intensity signal (arbitrary units, AU) along the DV axis of the neural tube (NT), expressed as the percentage of the NT length (bar plots: mean ± s.e.m). (C) Quantification of the number of cells positive for the indicated TFs (dots: values per transverse section, normalised to those of heterozygous littermates to account for stage variations across littermates, dot shapes: independent embryos; bar plots: mean ± s.e.m; Mann-Whitney U test: *p<0.05; **p<0.01; ***p<0.001; ****p<0.0001).

Among those TFs, we focused on PAX3 and PAX7, two paralogous TFs acting in the dorsal part of the spinal cord. PAX3 is first induced at embryonic day 8.0 (E8) in mice, forming a DV gradient spanning the roof plate, neural crest cells (NCCs), all dorsal progenitors, and, in the brachial region, extending ventrally to the ventral p0 progenitors and occasionally the p1 domain (Goulding et al. 1991; Moore et al. 2013; Zagorski et al. 2017; Kicheva et al. 2014). Within 12-24 hours, its expression becomes uniform and limited to the dorsal spinal cord (Figure 1A)(Goulding et al. 1991; Moore et al. 2013; Zagorski et al. 2017; Kicheva et al. 2014). At this stage, PAX7 is also induced, with lower levels in dp1 and dp2 cells and higher levels in dp3–dp6 cells (Figure 1A). While the absence of PAX7 does not cause significant spinal cord development defects, the loss of PAX3—and even more so in combination with PAX7—leads to DV patterning defects in the spinal cord (Gard et al. 2017; Mansouri and Gruss 1998; Mansouri et al. 1996; Nakazaki et al. 2008). The dp4 to dp6 dorsal progenitors give rise to ectopic ventral V0 and/or V1 INs in *Pax3^-/-^* embryos, with an even greater occurrence in *Pax3^-/-^; Pax7^-/-^* embryos (Gard et al. 2017). The fate of dp1 to dp3 progenitors in these mutants remains unexplored. However, the decreased expression of *Neurog2* and *Hes1*—two markers of these progenitors—in the absence of PAX3 suggests a role for PAX in their specification (Nakazaki et al. 2008; Ichi et al. 2011).

The transcriptional mechanisms by which PAX3 and PAX7 regulate the fate of dorsal progenitor pools remain poorly understood, but evidence highlights their versatility, with PAX proteins capable of both transcriptional activation and repression (Moore et al. 2013; Ichi et al. 2011; Nakazaki et al. 2008; Gard et al. 2017; Muhr et al. 2001). By analysing mutants where PAX3 activity is directed toward either repression or activation, we previously demonstrated that PAX proteins preserve the identities of dp4 to dp6 regions by inhibiting the expression of *Dbx1* and *Prdm12*, key determinants of the ventral p0 and p1 fates (Gard et al. 2017). While this repressive activity aligns with evidence that PAX7, like ventral spinal cord specification TFs, can bind *in vitro* to co-repressors Groucho/TLE (Muhr et al. 2001), it stands in contrast to their predominant role as transcriptional activators described in other developmental tissues (Mayran et al. 2015). In these tissues, PAX3 and PAX7 remodel chromatin promoting transcription through diverse mechanisms. PAX7 acts as a pioneer TF capable of binding and opening chromatin regions making them accessible to other TFs (Budry et al. 2012; Gouhier et al. 2024; Mayran et al. 2015). It also promotes enhancer-like epigenetic signatures at bound CRMs (Kawabe et al. 2012; McKinnell et al. 2008; Diao et al. 2012; Mayran et al. 2019), whereas PAX3 facilitates the establishment of three-dimensional chromatin structures, enhancing promoter-enhancer interactions (Magli et al. 2019). Such activator functions of PAX are likely to operate in the spinal cord. However, only a few enhancers whose activity depends on PAX3 have been identified near *Pax3*, *Neurog2*, and *Hes1* (Moore et al. 2013; Nakazaki et al. 2008; Ichi et al. 2011).

To investigate the transcriptional potential of PAX3 and PAX7 and their role in the DV patterning of spinal cord progenitor pools, we combined genetic and genomic approaches in mouse embryos and organoids. Our findings reveal that the transcriptional activity of PAX3 and PAX7 differs between dp1-dp3 and dp4-dp6 progenitors and depends on the CRMs they target. Across all dorsal progenitors, PAX proteins bind CRMs near ventral identity genes (p0 and p1), acting as repressors and contributing to H3K27me3 deposition. In dp1–dp3 progenitors exposed to BMPs, PAX-mediated repression coexists with transcriptional activation operating at distinct CRMs, promoting chromatin accessibility and driving H3K4me2 deposition and activating gene expression required for dI1-dI3 IN specification. Thus, the dual transcriptional role of PAX3/7, which expands the regulatory capacity of these TFs, constitutes a powerful mechanism for guiding the spatially organized diversification of cell types within the spinal cord.

## Results

### Spinal cellular diversity emerges and organizes through dorsoventrally modulated PAX3/7 transcriptional activity

To first investigate the spatial dynamics of PAX3 and PAX7 function as transcriptional activators in the embryonic spinal cord, we used the *P34::tk::nLacZ* reporter transgene (Relaix et al. 2003) (Figure 1B). This construct drives *LacZ* expression through a concatemer of five PAX3/7 binding sites recognised by PAX3 and PAX7 paired DNA-binding domain, located upstream of the *thymidine kinase* (*tk*) minimal promoter (Relaix et al. 2003). Reporter activity was assessed by β-galactosidase immunolabelling alongside GFP from a *Pax3GFP* knock-in allele in E9.5 mouse embryos, a stage when PAX3 and PAX7 are expressed and progenitors have distinct DV identities (Figure 1B). While PAX3-GFP was uniformly expressed within its domain at this stage, β-galactosidase activity showed a DV gradient, high dorsally and undetectable ventrally (Figure 1B). Co-immunolabelling of β-galactosidase and OLIG3, which marks the dp1-dp3 progenitor domain, showed that the reporter activity’s ventral boundary aligned with the dp1-dp3 domain boundary (Figure 1B). These results confirmed that in the embryonic spinal cord (Moore et al. 2013; Ichi et al. 2011; Nakazaki et al. 2008), PAX proteins activate gene expression specifically in dp1-dp3 but not in dp4-dp6 progenitors, indicating a context-dependent regulation of their transcriptional activity.

To assess the role of spatially modulated PAX3/7 activity in IN diversity (Figure 1C, S1), we first analysed the effects of PAX3 and/or PAX7 loss in mouse embryos carrying compound null knock-in alleles for *Pax3* (*Pax3^GFP^*) and/or *Pax7* (*Pax7^LacZ^*) (Figure 1C; S1A-C) (Mansouri et al. 1996; Zalc et al. 2015). We monitored IN subtypes specification at E11.5 using immunostaining for the TFs LHX2 (dI1), FOXD3 (dI2, V1), ISLET1/2 (dI3, pMN), LBX1 (dI4–dI6), LMX1B (dI5) and FOXP2 (high in V1) alongside GFP driven by *Pax3* locus (Figure 1A,C; S1).

The number and distribution of INs were the same in *Pax7^LacZ/LacZ^* embryos as in wild-type (WT) or double-heterozygous embryos (*Pax3^GFP/+^; Pax7^LacZ/+^*) (Figure 1C; S1A), consistent with the absence of a central nervous system phenotype in *Pax7* null mutants (Mansouri et al. 1996). In contrast, *Pax3^GFP/GFP^* embryos exhibited a reduced number of dI2 (FOXD3^+^), dI3 (ISLET1/2^+^), and dI5 (LMX1B^+^) INs, reaching between one-third and half the number observed in WT embryos (Figure 1C; S1A). The phenotype of *Pax3^GFP/GFP^; Pax7^LacZ/LacZ^* embryos was more severe, with a reduction of at least two-thirds in the number of all six types of dorsal INs (dI1 to dI6) (Figure 1C; S1A,B). Moreover, dorsal INs were mispositioned, as exemplified by dI4-6 (LBX1^+^) INs, which were found in the dorsal-most region of the spinal cord and extended into its dorsomedial part, a domain normally occupied exclusively by dI1-dI3 (LBX1^-^) INs (Figure S1A, arrows). This reduction in dorsal INs was accompanied by an increase in FOXD3^+^ and FOXP2^high^ V1 Ins (Gard et al. 2017; Mansouri and Gruss 1998) (Figure 1C, S1A,B). V1 INs arose not only from ventral progenitors PAX/GFP negative but also from dorsal PAX/GFP expressing progenitors, in a region that normally produces dI4-6 (LBX1^+^) INs (Figure 1C, S1A,B). This shows a switch of identity for some progenitors from dorsal to ventral. Collectively, these results highlight the partially redundant roles of PAX3 and PAX7 in the production of all dI1-dI6 INs and their spatial distribution along the DV axis of the spinal cord, with PAX3 playing a dominant role in dI1-dI3 IN production.

To determine which of the PAX3/7 activities—activator or repressor—is involved in generating dorsal spinal cord cellular diversity, we analysed the phenotypic consequences of modifying PAX3 transcriptional potential. We examined INs in E11.5 mouse embryos carrying conditional *Pax3* knock-in alleles expressing either an activator or repressor variant, with or without a *Pax3GFP* allele (Figure 1C, S1A,B) (Bajard et al. 2006; Relaix et al. 2003). These variants consisted of the DNA-binding domain of PAX3 fused to either the transactivation domain of FOXO1 (PAX3-FOXO1) or the repressor domain of Engrailed (PAX3-EnR) (Bajard et al. 2006; Relaix et al. 2003).

Strikingly, expressing the repressor PAX3-EnR in *Pax3*^+^ dorsal spinal progenitors permitted differentiation into dI4-dI6 INs while inhibiting dI1 to dI3 identities. In *Pax3^Pax3EnR/+^* E11.5 embryos, dI1 INs (LHX2^+^) and dI2 (FOXD3^+^) INs were reduced, whereas dI3-dI6 (ISLET1/2^+^, LBX1^+^, LMX1^+^) populations remained unchanged (Figure 1C, S1A). Notably, in *Pax3^Pax3EnR/GFP^* embryos, all dI1-dI3 INs were absent, and dI4-dI6 INs were generated, they were fewer and abnormally localized dorsally (Figure 1C, S1A). Conversely, the PAX3-FOXO1 activator shifted dorsal progenitor identities toward a V1-, dI1-, or dI2-like fate. At E11.5, *PAX3^PAX3-FOXO1/+^* embryos, exhibited fewer dI4-dI6 (LBX1^+^) INs and dI5 (LMX1B^+^) INs than WT embryos (Figure 1C, S1A). Additionally, in the dorsal-most region, dI3 (ISLET1/2+) IN production was reduced, while dI1 (LHX2+) and dI2 (FOXD3+) INs were more abundant and extended ventrally compared to controls (Figure 1C; S1A,B). Moreover, ectopic FOXD3^+^ neurons emerged from the ventricular (progenitor) region, from where LBX1^+^ dI4-dI6 INs typically arise. Among these, some expressed high levels of FOXP2, marking them as V1 INs, while others, lacking or expressing low FOXP2, resembled dI2-like cells (Figure S1B).

In summary, a forced PAX repressive activity impairs dI1-dI3 IN production, while a forced activation partially skews dp4-dp6 progenitors toward a V1- or dI2-like fate. Hence, the balance between the activator and repressor activities of PAX3 and PAX7, regulated along the DV axis, is crucial for shaping the generation of dorsal interneuron subtypes. This underscores the importance of identifying the *cis*-regulatory elements through which these TFs fine-tune distinct transcriptional programs along the DV axis of the spinal cord.

### mESC-derived organoid models unveil spinal IN specification through dual PAX3/7 transcriptional dynamics

To dissect PAX3/7 dual activity in dorsal progenitors, we generated mESC-derived spinal organoids, overcoming the challenge of accessing limited progenitor pools in embryos while recapitulating their differentiation dynamics (Duval et al. 2019). Spinal organoids transitioned from pluripotency (*Klf4^+^; Fgf5^+^; Pou5f1^-^*) to an epiblast state (*Klf4^-^; Fgf5^+^; Pou5f1^+^*) within 48h, then to a caudal neural state (*Nkx1.2^+^; Cdx2^+^; Sox1^low^; Pax6^low^*) by 66h (Figure S2A). By day 3, *Sox1* and *Pax6* increased as *Nkx1.2* and *Cdx2* decreased, with *Tubb3*^+^ neurons emerging by day 6 and peaking at day 7 (Figure S2A). DV identities were modulated by BMP4 exposure between days 3-4 (Figure S2A,B). Untreated organoids (ø) contained mixed dp4-dp6 and ventral p0/p1 progenitors (*Pax3/7^+^; Olig3^-^* dp4-dp6*; Prdm12^+^* p1; *Dbx1^+^* p0, dp6; *Lbx1^+^* dI4-dI6), while BMP4 favoured dp1-dp3 fates progenitors (*Pax3/7^+^; Olig3^+^* dp1-dp3; *Pou4f1^+^* dI1-dI3, dI5). PAX3 and PAX7 were detected from day 3, but their expression varied: BMP4 increased PAX3 while slightly reducing PAX7 (Figure 2B, S2B). The *P34::tk::nLacZ* transgene confirmed PAX3/7 transcriptional activation in BMP4-treated OLIG3^+^ progenitors, mirroring its dorsal restriction to dp1 and dp3 progenitors in developing embryos (Figure 1B, 2C).

**Figure 2:**
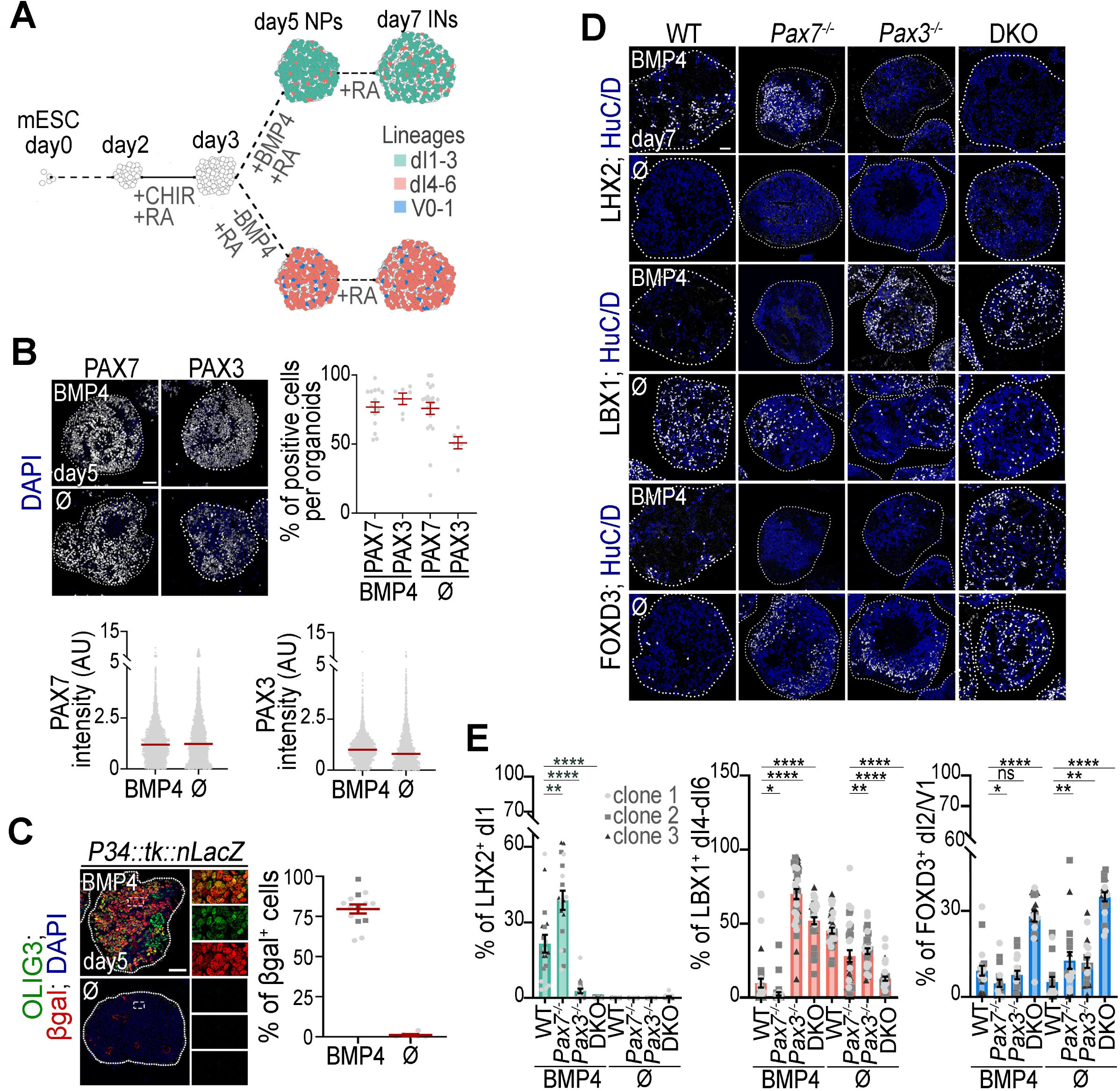
Expression, activity, and function of PAX3 and PAX7 during spinal neuronal diversity generation in organoids treated or not with BMP4. **(A)** Protocol for generating spinal organoids from mESCs, enriched in dp4-dp6 progenitors (BMP4-free, ø condition) or dp1-dp3 progenitors (with BMP4) at day 5, and their associated INs at day 7 of differentiation. Few ventral cells of the V0 and V1 lineages are present in organoids untreated with BMP4. **(B)** Immunostaining for PAX7 or PAX3 (white) and DAPI-stained nuclei (blue) on sections of day 5 WT spinal organoids treated or not with BMP4. Quantification of the percentage of PAX7^+^ or PAX3^+^ cells per organoid (dots: individual organoid values; bar plots: mean ± s.e.m.) or PAX7 or PAX3 intensity levels (dots: individual nuclear values in arbitrary units (AU); bar plots: mean ± s.e.m.). **(C) Images:** Immunostaining for β-galactosidase (βgal; red), OLIG3 (green) and DAPI-stained nuclei (blue) on sections of day 5 *P34::tk::LacZ* spinal organoids treated or not with BMP4. Right panels are magnified views of the squares shown in the left panels. **Graphs:** Quantification of the percentage of βgal^+^ cells per day 5 organoids treated or not with BMP4, (dots: organoids (left graph) and bar plots: mean ± s.e.m.). **(D)** Immunostaining for LHX2, LBX1, FOXD3 (white) and DAPI stained nuclei (blue) on sections of day 7 spinal organoids with the indicated genotypes, treated or not BMP4. **(E)** Quantification of the percentage of cells expressing LHX2, LBX1, FOXD3 within day7 organoids with the indicated genotypes (dots: values per transverse section, dots shapes: independent clones; bar plots: mean ± s.e.m.; Mann-Whitney U test: *p<0.05; **p<0.01; ***p<0.001; ****p<0.0001). In all images, dotted lines delineate the spinal cord, and scale bars represent 50 μm. WT: Wild-type; DKO: *Pax3; Pax7* double mutant.

Using CRISPR, we generated three independent *Pax3^-/-^, Pax7^-/-^*, and *Pax3; Pax7* double knockout (DKO) mESC lines, by excising the first two exons and part of the third exon of the *Pax3* and/or *Pax7* genes. Immunostaining confirmed the absence of both proteins in day 5 mutant organoids (Figure S2C). In all mutant organoids, neural commitment remained intact, with SOX1^+^ cells exceeding 96% at day5 (Figure S2E). The unreduced number of HuC/D^+^ neurons at day7 indicating that terminal differentiation remained unaffected in absence of the PAX (Figure S2F). However, IN composition was altered, as revealed by LHX2 (dI1), LBX1 (dI4-dI6), FOXD3 (dI2, V1), POU4F1 (dI1-3, dI5), and PAX2 (dI4, dI6, V0, V1) immunostaining (Figure 2D,E; S3).

In BMP4-treated organoids, the loss of PAX7 had minimal impact on cell composition. Similar to controls, *Pax7^-/-^* organoids contained over 30% LHX2^+^ dI1 INs and fewer than 7% LBX1^+^ dI4-dI6 cells (Figure 2D, E). In contrast, *Pax3^-/-^* organoids exhibited a sixfold reduction in LHX2^+^ dI1 INs, while LBX1^+^ dI4-dI6 INs increased, comprising nearly 75% of HuC/D^+^ neurons versus < 10% in controls (Figure 2D,E). The combined loss of PAX3 and PAX7 (DKO) further exacerbated the ventralisation of cell identities. LHX2 expression was nearly absent, while LBX1^+^ dI4-dI6 INs made up half of HuC/D^+^ neurons. Additionally, 30% of neurons were FOXD3^+^ (Figure 2D,F), but lacked POU4F1, indicating a V1-like rather than dI2-like IN identity (Figure S3A). The presence of PAX2^+^; LBX1^-^ V0 and V1-like INs in DKO, absent in WT organoids further confirmed this ventral identity shift (Figure S3B). To determine whether the ventralisation observed in BMP4-treated *Pax3^-/-^* and DKO organoids resulted from impaired BMP4 response—essential for dI1-dI3 specification (Tozer et al. 2013; Zechner et al. 2007; Duval et al. 2019; Le Dréau et al. 2012; Andrews et al. 2017)—we analysed phosphorylated SMAD1/5/9, the transcriptional effectors of BMP signalling. One hour post-BMP exposure, when signalling peaks (Duval et al. 2019), phosphorylated SMAD levels at the organoid periphery were comparable across all genotypes (Figure S3C), ruling out BMP signalling defects as the cause of ventralisation in *Pax3^-/-^* and DKO organoids. Even without BMP4, loss of PAX3 or PAX7 loss, and especially their combined loss, led to ventralisation of neuronal identities (Figure 2D,E; S3A,B). LBX1^+^ dI4-dI6 INs decreased from 50% in controls to 30% in single mutants, and below 10% in DKO organoids (Figure 2D,E). Conversely, V0 and V1 INs (FOXD3^+^, PAX2^+^, POU4F1^-^) increased from ∼10% in controls to over 15% in single mutants and up to 30% in DKO organoids (Figure 2D,E; S3A,B).

In conclusion, as in embryos, PAX3/7 transcriptional activation in organoids is restricted to dp1-dp3 progenitors and absent in dp4-dp6. In both models, PAX3 and PAX7 cooperate to prevent ventral identity acquisition, with PAX3 playing a dominant role in dp1-dp3. However, unlike *in vivo*, PAX7 loss significantly impacts dI4-dI6 IN production in organoids. This discrepancy may stem from reduced PAX3 expression in dp4-dp6 cells *in vitro*, likely due to the need for finer BMP and Wnt signalling modulation during differentiation (Moore et al. 2013; Alvarez-Medina et al. 2008; Timmer et al. 2005).

### PAX3 and PAX7 differentially shape dorsoventral gene regulatory networks

To investigate PAX3/7-regulated transcriptional programs in dp1-dp3 and dp4-dp6 progenitors, RNA-seq was performed on WT, single-mutant, and DKO organoids (Figure 3A-D). Using three independent cell lines per genotype, organoids were treated or untreated with BMP4 and harvested at day 6, when progenitors and INs coexist (Figure 3; Table S1).

**Figure 3.**
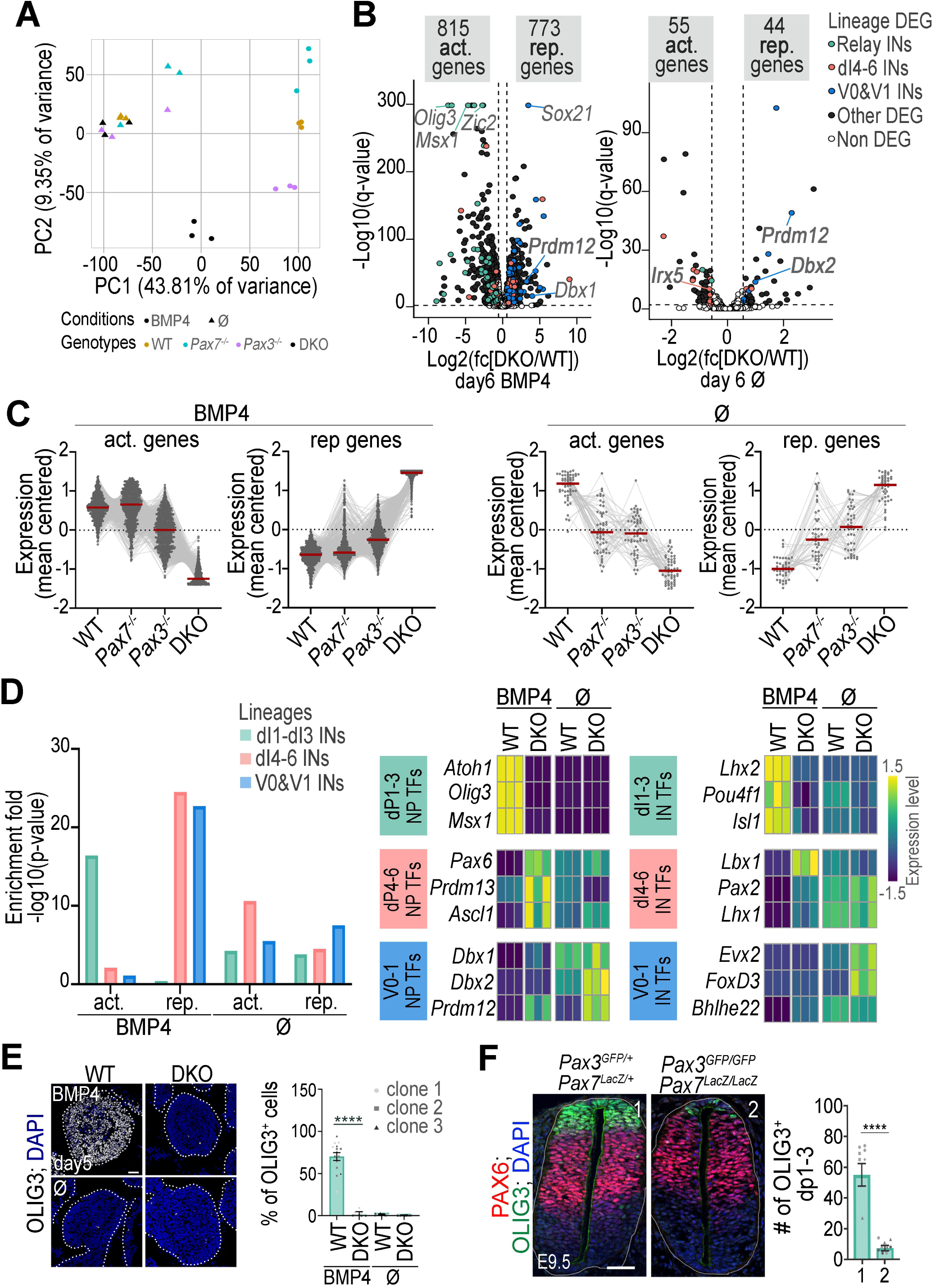
Transcriptional programmes regulated by PAX3 and PAX7 activity in organoids treated or not to BMP4. **(A)** Principal component analysis of day 6 organoids with the specified genotype, treated or not with BMP4. **(B)** Volcano plots of comparing gene expression between WT and DKO day 6 organoids, treated or not with BMP4. The y-axis shows statistical significance (-log10(q-value)) and the x-axis represents log2(fold change). White dots: non-significant differentially expressed genes (DEG), black/coloured dots: significant DEGs. Blue, pink, and green dots highlight master TFs for ventral, dI1-dI3, and dI4-6 IN lineages, respectively. **(C)** Normalised mean-centred expression levels of genes downregulated or upregulated in DKO vs. WT day6 organoids, treated or not with BMP4, shown for WT, *Pax7^-/-^*, *Pax3^-/-^*, and DKO organoids (dots/lines: individual genes; bars: mean ± s.e.m.; n=3 clones/genotype). **(D) Left:** Enrichment of genes encoding dI1-dI3, dI4-6, or V0-V1 specifiers in the list of genes downregulated (down.) or upregulated (up.) in DKO organoids compared to WT day 6 organoids, with or without BMP4. **Right:** Heatmaps showing normalised mean-centred mRNA expression of TFs marking progenitors (NPs) or neurons (IN) in the dI1-dI3, dI4-6 or V0/V1 INs lineages assessed by RNA-seq in day 6 WT or DKO organoids, treated or not with BMP4, in 3 independent experiments. Fold induction across samples are colour-coded in blue (lower levels) to yellow (higher levels). **(E)** Immunostaining for OLIG3 (white) and DAPI-stained nuclei (blue) on sections of day 5 spinal organoids with the indicated genotype, treated or not with BMP4. Quantification of the percentage of OLIG3^+^ cells in these organoids (dots: values per transverse section, dots shapes: independent clones; bar plots: mean ± s.e.m; Mann-Whitney U test: ***p<0.001; ****p<0.0001; ns: non-significant). **(F)** Immunostaining for OLIG3 (green), PAX6 (red) and DAPI stained nuclei (blue) on transverse sections of E9.5 mouse embryos with the indicated genotypes and quantification of the number of OLIG3^+^ cells in such embryos (dots: values per transverse section, dots shapes: independent embryos; bar plots: mean ± s.e.m; Mann-Whitney U test: ****: p<0.0001).

Principal component analysis showed transcriptomic profiles primarily segregated by BMP4 treatment, with genotype-based segregation evident only in BMP4-treated organoids (Figure 3A). Consistently, DESeq2 identified more differentially expressed genes (DEGs) in BMP4-treated *Pax3*/*Pax7* mutants than in untreated conditions (Figure 3B, Table S1). These DEGs included both PAX-activated (downregulated in DKO) and PAX-repressed (upregulated in DKO) genes, highlighting a stronger PAX3/7 impact on dp1-dp3 (BMP4) than dp4-dp6 (Ø) progenitors. The number of DEGs was highest in DKO organoids, exceeding those in single mutants (Table S1). Without BMP4, gene expression in single mutants was intermediate between WT and DKO, suggesting overlapping PAX3 and PAX7 associated gene networks (Figure 3C). Under BMP4, genes deregulated in DKO were partially affected in *Pax3^-/-^*but unchanged in *Pax7^-/-^* mutants, confirming PAX3 as the primary transcriptional regulator in BMP4-treated conditions (Figure 3C)

To identify PAX3/7 effectors, we defined gene signatures for dp1-dp3, dp4-dp6, p0-p1 progenitors as well as their associated INs, using single-cell RNA-seq and histological data from embryonic mouse and chicken spinal cords (Rekler et al. 2024; Sagner and Briscoe 2019; Delile et al. 2019). We then analysed the enrichment for these molecular signatures among DEGs upon Pax3/Pax7 loss (Figure 3D; Table S1). In BMP4-treated organoids, PAX3/7-activated genes were highly enriched in dp1-dp3/dI1-dI3 signatures (Figure 3D), including key TFs for specification and differentiation, such as *Olig3* (essential for dI2/dI3 INs (Müller et al. 2005)), *Atoh1*, and *Msx1* (required for dI1 INs (Gowan et al. 2001; Duval et al. 2014)) (Table S1), as well as IN markers *Lhx2*, *Pou4f1*, and *Isl1* (Sagner & Briscoe 2019). To validate this, we assessed OLIG3 distribution in DKO day 5 organoids and E9.5 mouse embryos (Figure 3E). OLIG3 was detected in over 60% of BMP4-treated WT organoid cells but was absent in untreated WT organoids (Figure 3E). In DKO BMP4-treated organoids, OLIG3 induction was nearly abolished (Figure 3E), mirroring a sixfold reduction in OLIG3^+^ cells in E9.5 *Pax3; Pax7* double mutant embryos (Figure 3F). Conversely, PAX3/7-repressed genes in BMP4-treated organoids were enriched in dp4-dp6/dI4-dI6 and p0-p1/V0-V1 signatures. These included *Pax6* (highly expressed in intermediate spinal cord regions (Briscoe et al. 2000)), *Prdm13* and *Ascl1* (which drive dI4-dI6 INs (Helms et al. 2005; Chang et al. 2013)), and IN markers *Lbx1*, *Pax2*, and *Lhx1* (dI4-dI6) (Sagner & Briscoe 2019). Additionally, p0/p1 regulators *Prdm12*, *Dbx1, Dbx2* (Thélie et al. 2015; Pierani et al. 2001) and V0/V1 IN markers *Foxd3, bHlhe22*, and *Evx2* were also repressed (Sagner & Briscoe 2019). In untreated organoids, genes deregulated in the absence of PAX factors are also enriched in those involved in the specification of dp1-dp3, dp4-dp6, and d0-p1 lineages, though these enrichments are less pronounced than in BMP4-treated conditions (Figure 3D). Notably, p0-p1 and V0/V1 markers were more strongly enriched among PAX-repressed genes, whereas dp4-dp6 lineage markers were predominantly found among PAX-induced genes (Figure 3D).

These findings demonstrate that PAX3 and PAX7 orchestrate transcriptional programs critical for dp1-dp3, dp4-dp6, and p0-p1 specification, explaining the altered IN fates observed in their absence. PAX factors universally repress p0-p1 programs, preventing dorsal progenitors from acquiring V0/V1 identities. However, dp1-dp3 programs, exclusive to BMP4 conditions, rely entirely on PAX activation. In contrast, PAX3/7 promote dp4-dp6 programs without BMP4 but repress them in its presence, reflecting BMP-dependent modulation of PAX-regulated networks.

### PAX3/7 drive distinct chromatin state dynamics along the DV axis

To uncover CRMs driving PAX3/7 transcriptional effects, we generated mESC lines with 3xFLAG-tagged *Pax3* or *Pax7*, enabling CUT&Tag-based profiling of genomic recruitment sites in day 5 organoids (Figure 4A, Table S2). We identified 3,297 PAX3/7-enriched peaks, primarily in intronic and distal intergenic regions (Figure 4A, Table S2; Figure S4A). Using GREAT (McLean et al. 2010), we mapped nearby genes, which were significantly enriched among PAX3/7-activated and repressed targets in transcriptome analyses, suggesting these regions act as CRMs, promoting or inhibiting gene expression (Figure S4B).

**Figure 4.**
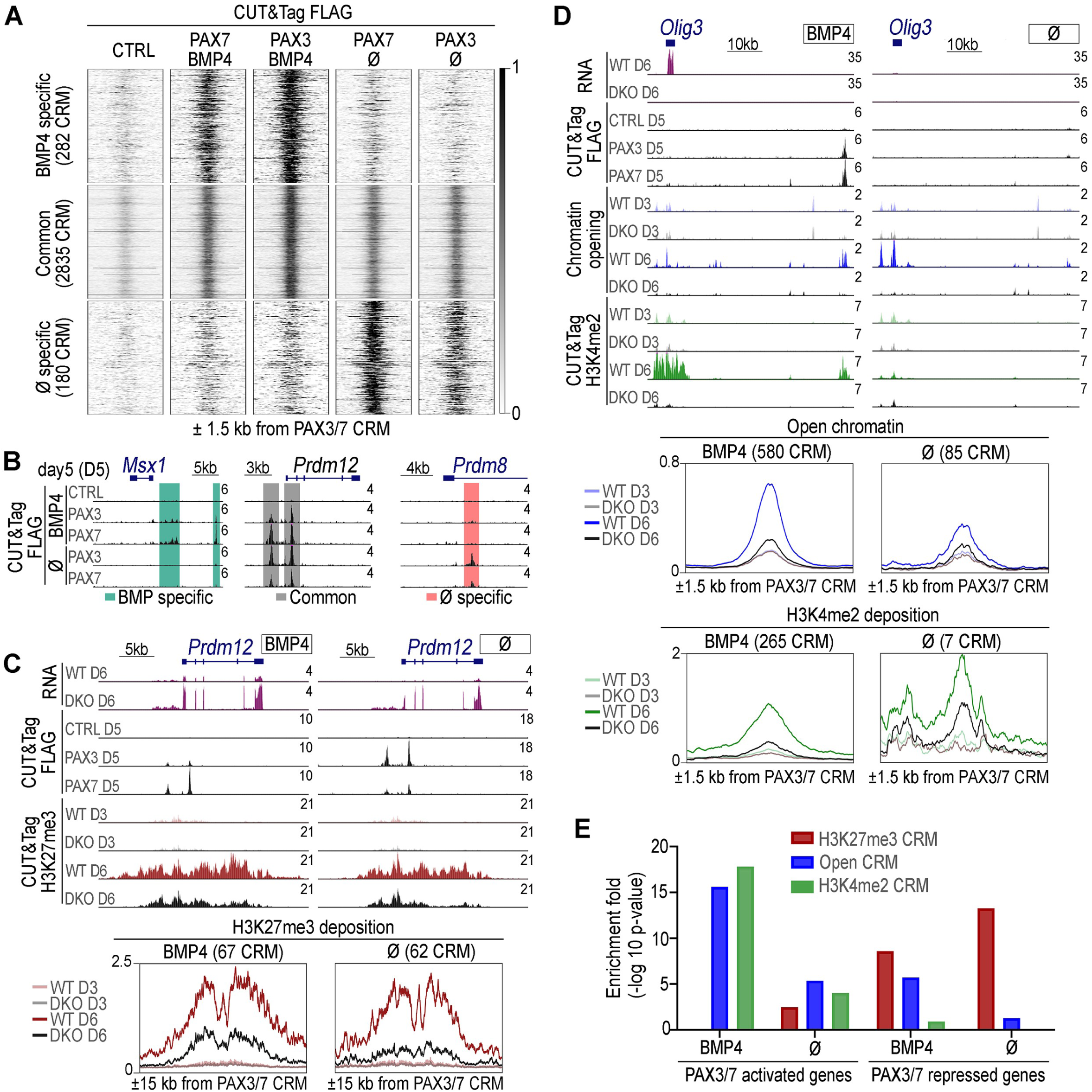
Dynamics of chromatin states at PAX3/7-bound CRMs after their loss and BMP4 exposure. **(A)** Heatmaps of normalised FLAG CUT&Tag signals for all PAX3/7-bound CRMs identified in organoids expressing FLAG-PAX3 (PAX3), FLAG-PAX7 (PAX7), or not (CTRL), treated or untreated with BMP4 at day 5 of differentiation, over a 1.5 kb region upstream and downstream of the CRM centres (n = 3297). Three groups of PAX3/7-bound CRMs are indicated: CRMs specific to BMP4-treated organoids, those specific to untreated organoids, and those shared between both conditions. **(B)** UCSC Genome Browser screenshots showing normalised FLAG CUT&Tag read distributions in wild-type (CTRL) and FLAG-PAX3 (PAX3) or FLAG-PAX7 (PAX7) organoids, with or without BMP4 treatment, at day 5 of differentiation. Scales are in CPM (counts per million). Coordinates: chr5:37816721-37837786 (left, scale bar: 5 kb) and chr7:49629789-49642743 (right, scale bar: 2 kb). Common recruitment sites are highlighted in grey, BMP4-specific in green, and untreated-specific in salmon-pink. **(C)** UCSC Genome Browser screenshots showing normalised RNA-seq (RNA), FLAG, or H3K27me3 CUT&Tag read distributions in wild-type (CTRL), FLAG-PAX3 (PAX3) or FLAG-PAX7 (PAX7) expressing, and *Pax3; Pax7* double-knockout (DKO) organoids, at days 3, 5 or 6 (D3, D5, D6) of differentiation, treated or not BMP4. Scales are shown in CPM. Coordinate: chr2:31627589– 31663216, scale bar: 5kb. Metaplots at the bottom show H3K27me3 enrichment over a 15 kb region upstream and downstream of the centres of PAX3/7-bound CRMs displaying significant differential H3K27me3 deposition between CTL and DKO organoids. Data are shown for WT organoids at days 3 and 6, and DKO organoids at day 6 of differentiation. **(D)** UCSC Genome Browser screenshots showing normalised RNA-seq (RNA), FLAG CUT&Tag, ATAC-seq (chromatin accessibility), or H3K4me2 CUT&Tag read distributions in wild-type (WT), FLAG-PAX3 (PAX3), FLAG-PAX7 (PAX7) expressing, and Pax3; Pax7 double-knockout (DKO) organoids, treated or untreated with BMP4, at days 3, 5, or 6 (D3, D5, D6) of differentiation. Scales are shown in CPM. Coordinates: chr10:19355317-19416668, scale bar: 10 kb. Metaplots at the bottom show ATAC-seq and H3K4me2 CUT&Tag signal enrichment over a 1.5 kb region upstream and downstream of the centres of PAX3/7-bound CRMs displaying significant differential ATAC-seq signal or H3K4me2 deposition between CTRL and DKO organoids at day 6 of differentiation. Data are shown for WT organoids at days 3 and 6, and for DKO organoids at day 6 of differentiation. **(E)** Enrichment of genes activated or repressed by PAX3 and PAX7 (identified as downregulated and upregulated genes in Figure 3, respectively) among genes located near PAX3/7-bound CRMs displaying significant differential ATAC-seq signals (blue), H3K4me2 deposition (green), or H3K27me3 deposition (dark red) between CTL and DKO organoids at day 6 of differentiation (bars: -log10(p-value)).

PAX3 and PAX7 showed similar recruitment patterns, with peak intensities reflecting their expression: PAX3 peaks were stronger in BMP4-treated organoids, while PAX7 peaks stronger in untreated ones (Figure 4A). However, recruitment sites differed significantly with BMP4 treatment, highlighting context-dependent binding (Figures 4A,B). Of 3,297 PAX3/7-bound CRMs, 2,835 were shared, including two within *Prdm12* locus (p1 marker) (Figures 4A,B). BMP4-treated organoids had 282 unique CRMs, such as those near *Msx1* (dp1-dp3), while 180 CRMs were exclusive to untreated organoids, including intronic regions of *Prdm8* (ventral marker)(Figures 4A,B).

To assess whether PAX3/7 recruitment is influenced by CRM motif composition, we performed motif enrichment analyses across three CRM categories: BMP4-specific, shared, and untreated-specific (Figure S4C,D). First, we examined motifs recognized by PAX DNA-binding domains—the Paired domain and Homeodomain (Blake and Ziman 2014). These included motifs bound by the Paired domain (PrD), a Homeodomain dimer (HD-HD), or both domains simultaneously (PrD-HD) (Figure S4C)(Mayran et al. 2015). All three motifs were enriched in shared CRMs, with PrD-HD predominating. BMP4-specific CRMs favoured PrD-HD and PrD motifs, while untreated-specific CRMs showed lower PAX motif representation, suggesting indirect recruitment (Figure S4C).

We then analysed motif enrichment for other TFs in PAX3/7-bound CRMs (Figure S4D). Motifs recognized by key regulators of embryonic neural cells—such as homeodomain, zinc-finger-containing (SP/KLF), and SOX TFs—were enriched across all PAX CRM categories, suggesting that PAX3 and PAX7 cooperate with these factors in neural specification, as observed for other specification TFs (Zhao et al. 2022; Bailey et al. 2006; Oosterveen et al. 2012, 2013; Nishi et al. 2015; Kutejova et al. 2016; Chang et al. 2013; Borromeo et al. 2014; Lodato et al. 2013). Additionally, motifs for ZIC TFs were specifically enriched in BMP4-specific CRMs (Figure S4D). Given the dorsal restriction of ZIC TFs in the spinal cord (Inoue et al. 2004), such PAX-ZIC interactions may facilitate locus-specific recruitment in BMP4-treated organoids.

We then characterized the chromatin state of PAX3/7-bound CRMs during differentiation and assessed whether this state depended on PAX activity. We performed CUT&Tag profiling mapped H3K27me3 (Polycomb-mediated repression) and H3K4me2 (MLL-associated activation) (Ernst et al. 2011; Wang et al. 2014) in control and mutant organoids. Additionally, ATAC-seq identified accessible genomic regions (Figure 4C,D).

Among PAX3/7-bound CRMs, ∼100 were marked by H3K27me3 in both BMP4-treated and untreated organoids, with half shared between conditions (Figure 4C, S4E). This mark was dynamically deposited during differentiation—absent at day 3, detectable by day 5, and more pronounced by day 6 (Figure 4C, S4E). Notably, H3K27me3 levels dropped at over 60 of these CRMs in absence of PAX3 and PAX7, regardless of BMP4 treatment (Figure 4C). Nearby genes were enriched for PAX3/7-repressed targets, reinforcing their role in transcriptional repression (Figure 4E). Several CRMs near *Prdm12* gene illustrated this PAX-dependent regulation (Figure 4C). A partial reduction in H3K27me3 levels was observed in *Pax3^-/-^* or *Pax7^-/-^* organoids but was far more pronounced in DKO mutants (Figure S5B,C).

Unlike H3K27me3 deposition, which was restricted to a small subset of PAX3/7-bound CRMs, chromatin opening (ATAC-seq) was observed over half of these CRMs, regardless of culture conditions (Figure S4E). Some of these regions required PAX activity for opening (Figure 4D, S4E), with 580 CRMs opening in BMP4-treated organoids but only 85 in untreated ones (Figure 4D), indicating a strong BMP-dependent effect. Among BMP4-responsive CRMs, PAX3/7 opened both BMP4-specific and shared CRMs in nearly equal numbers (Figure S4E). However, 70% of BMP4-specific CRMs required PAX, compared to just 14% of shared CRMs (Figure S4E). More than half of PAX-dependent open regions were initially closed at day 3 but progressively opened during differentiation (Figure 4D, S4E, S5A), suggesting that PAX factors act as chromatin remodelers, similar to pioneer factors in the pituitary (Mayran et al. 2018).

Loss of PAX3 or PAX7 alone had milder effects on chromatin opening than their combined loss (Figure S5B,C). Additionally, 272 PAX3/7-bound CRMs were marked with H3K4me2, almost exclusively in BMP4-treated organoids (265), with deposition largely dependent on PAX activity (Figure 4D, S4E). Genes near open or H3K4me2-marked regions were predominantly PAX-regulated, reinforcing the role of these CRMs in PAX-mediated gene activation (Figure 4E).

These findings underscore the pivotal role of PAX proteins in shaping the chromatin landscape of CRMs in dorsal spinal progenitors. They regulate H3K27me3 deposition near repressed genes, particularly in dp1-dp3 and dp4-dp6 progenitors. In dp1-dp3 progenitors, PAX3 and PAX7 also promote chromatin accessibility at CRMs near activated genes, opening closed regions, maintaining open states, and facilitating enhancer-associated activation marks. Thus, PAX proteins exhibit dual transcriptional activity, modulated by progenitor identity and target CRM sequences.

### PAX3/7-bound silencers and enhancers shape DV gene expression restriction

We finally investigated the implication PAX3/7-bound CRMs in DV patterning of the spinal cord (Figure 5). First, we analysed the enrichment of DV patterning genes near PAX3/7-bound CRMs whose regulatory state depended on PAX activity (Figure 5A). PAX3/7-bound CRMs were classified into three categories based on PAX-dependent modifications: those marked by H3K27me3 deposition, H3K4me2 deposition, or chromatin opening. Genes located near opened and/or H4K4me2-marked CRMs were enriched for dp1-dp3 regulators, but only in BMP4-treated organoids (Figure 5A). Conversely, genes near H3K27me3 marked CRMs were strongly associated with ventral V0 and V1 IN fate, regardless of BMP4 treatment status (Figure 5A). Notably, dI4-dI6 regulators showed no significant enrichment near any of the PAX3/7-bound CRMs. These findings highlight a functional segregation of PAX dual activity: repression targets ventral gene CRMs, while activation acts on dp1-dp3 CRMs.

**Figure 5.**
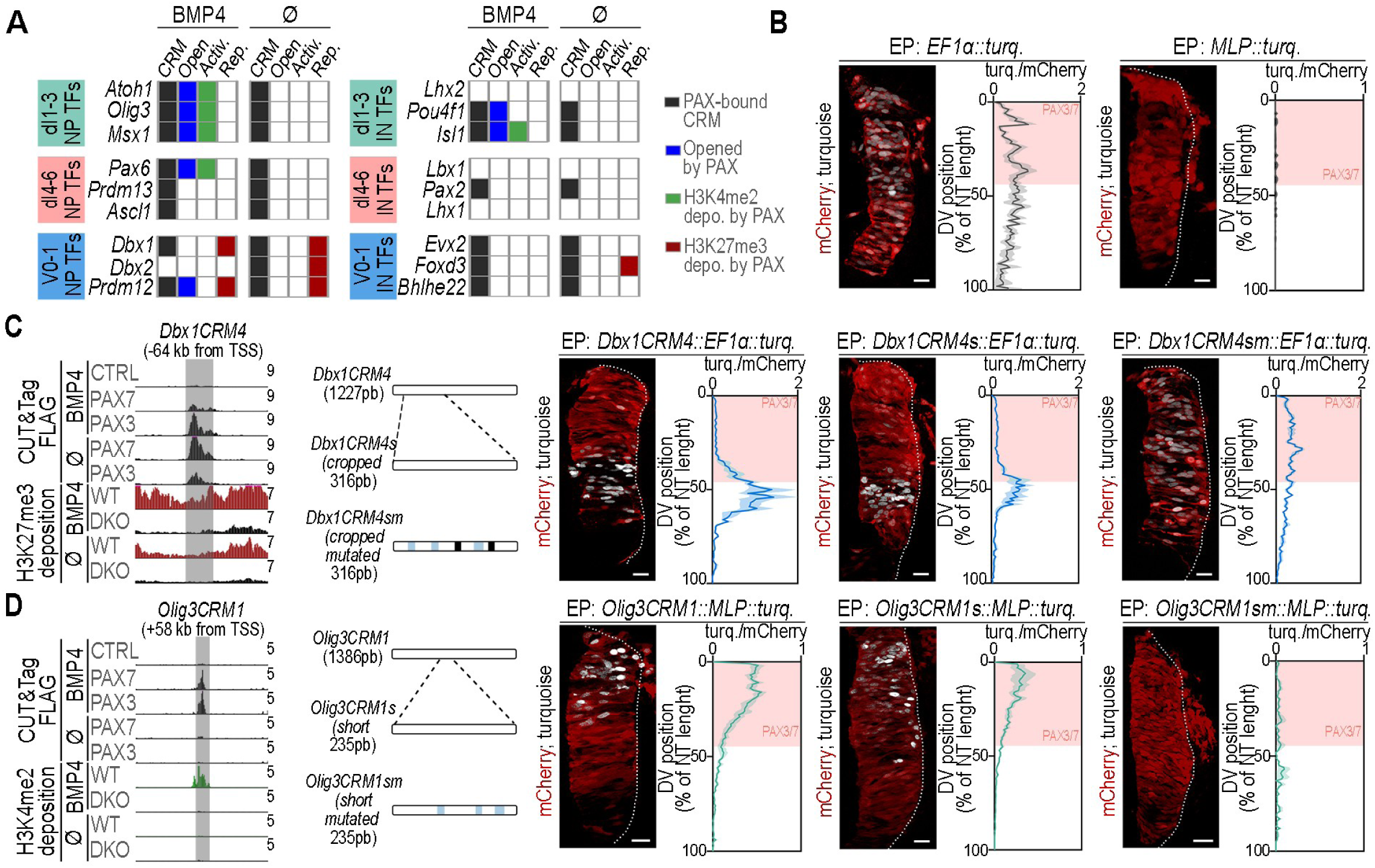
Silencer or enhancer activity of PAX3/7-bound CRMs nearby DV patterning genes. **(A)** Table indicating the presence of PAX3/7-bound CRMs (black) near DV patterning TFs driving relay, dI4-6, and V0/V1 INs differentiation, along with their chromatin state: opened by PAX (Open, blue), PAX-mediated H3K4me2 deposition (Activ. green), or PAX-mediated H3K27me3 deposition (Rep. dark red). **(B)** Fluorescence of mCherry (red) and Turquoise (white) on transverse sections of the chick neural tube, 24 hours post-electroporation with the reporter plasmids from *EF1α* or *MLP* promoters’ activity. Scale bars: 50µm. **(C,D) Left panels**: UCSC Genome Browser screenshots showing normalised FLAG (black tracks), H3K27me3 (dark red tracks) or H3K4me2 (green tracks) CUT&Tag read distributions in control (CTRL), FLAG-PAX3 (PAX3), FLAG-PAX7 (PAX7) at day 5 of differentiation and wild type (WT) and Pax3; Pax7 double-knockout (DKO) organoids at day 6 of differentiation, either treated or not BMP4. Read scales are shown in CPM. Grey bands highlight the position of *Dbx1CRM4* chr7:49697106-49701929 (**C**)) and *Olig3CRM1* CRMs (chr10:19412350-19416446 (**D**)). **Middle panels:** Position of the shortened (s), or shortened and mutated (sm) version of either *Dbx1CRM4* and *Olig3CRM1*, with the following colour scheme: black for PrD, lightblue for PrD-HD. **Right panels:** Fluorescence of mCherry (red) and Turquoise (white) on transverse sections of the chick neural tube, 24 hours post-electroporation with the indicated constructs. Scale bars: 50µm. Quantification of the ratio between Turquoise and mCherry signal intensities along the dorsoventral (DV) axis of the neural tube (NT) expressed as a percentage of NT length (bar plots represent mean ± s.e.m.; n≥4 embryos). Salmon-colored rectangles indicate the positions of PAX3/7-expressing progenitors.

We then assessed the transcriptional potential of selected CRMs and their spatial activity along the DV axis of the neural tube in chick embryos (Figure 5B-D, S6). Ten PAX-dependent H3K27me3-marked CRMs were cloned upstream of the *EF1α* promoter, which normally drives uniform *turquoise* transgene expression along the DV axis (Figure 5B). Among them, *Dbx1CRM4,* located near *Dbx1*, repressed *EF1α* promoter activity throughout the PAX3/7 domain, except in a ventral row likely corresponding to dp6 progenitors (Figure 5C, S6A). This indicates that *Dbx1CRM4* silences transcription in dp1-dp5 progenitors. To further investigate PAX involvement in this silencing activity, we shortened *Dbx1CRM4* and mutated its inner PAX binding sites (Figure 5C, Table S3). The shortened *Dbx1CRM4* (*Dbx1CRM4*s) retained its ability to repress *EF1α* promoter activity dorsally (Figure 5C). However, its mutated version (*Dbx1CRM4sm*) failed to do so, resulting in Turquoise expression along the neural tube without any DV restriction (Figure 5C).

Twenty PAX3/7-bound CRMs requiring PAX for chromatin opening and H3K4me2 deposition were cloned upstream of the minimal promoter (*MLP*), which is transcriptionally inactive on its own (Figure 5B). One-third of these CRMs activated *MLP* promoter activity in PAX3/7^+^ progenitors, including CRMs in the vicinity of *Olig3 (Olig3CRM1)*, *Msx1* (*Msx1CRM3*), and *Slit1* (*Slit1CRM1*)(Figure 5D, S6B,C). Strikingly, their enhancer activity was restricted to the most dorsal region of PAX3/7^+^ domain, corresponding to dp1 to dp3 progenitors, indicating spatially confined activation (Figure 5D, S6A-C). Shortened versions of these CRMs retained *MLP* activation, although some of them were extended ventrally, suggesting the loss of sequences restricting DV boundaries (Figure 5D, S6B,C). Furthermore, mutating PAX motifs in these truncated CRMs completely abolished transcriptional activity (Figure 5D, S6B,C).

These findings demonstrate that PAX-modulated CRMs function as silencers or enhancers, with distinct regulatory roles along the spinal cord DV axis. PAX3/7-bound enhancers are active specifically in dp1–dp3 progenitors, mirroring the *P34::tk::LacZ* reporter gradient (Figure 5C, 1B). In contrast, silencing extends more broadly within the PAX domain, coexisting with activation in dp1–dp3 progenitors.

## Discussion

Our study reveals that the paralogous TFs PAX3 and PAX7 play a pivotal role in generating spinal cord cellular diversity by guiding dorsal spinal progenitors (dp1 to dp6) along distinct differentiation trajectories. Their dual transcriptional activity—encompassing both activation and repression—is central to this process. Key parameters underlying the implementation of this dual activity were identified as critical for spinal cell fate patterning. First, like other TFs, such as FOXD3 in neural crest cells (NCCs) or reprogramming factors OCT4, SOX2, and KLF4 (OSK)(Katsuda et al. 2024; Lukoseviciute et al. 2018), PAX3/7-mediated activation and repression govern distinct differentiation programs: repression restricts ventral fates (p0-p1), while activation drives dorsal progenitor (dp1-dp3) differentiation. This segregation occurs at the genomic level, where PAX3/7-regulated enhancers and silencers occupy distinct genomic regions. Second, unlike FOXD3 and reprogramming factors—whose repressive and activating functions are temporally separated (Katsuda et al. 2024; Lukoseviciute et al. 2018)—PAX3/7 activities can coexist within the same cell type. FOXD3, for instance, first maintains NCC multipotency before shifting to a repressive role that commits cells to differentiation (Lukoseviciute et al. 2018). In contrast, PAX3/7-mediated activation and repression operate simultaneously within dp1–dp3 progenitors, as demonstrated by the co-occurrence of *Dbx1CRM4* silencing and *Olig3CRM1* enhancer activity (Figure 5). Finally, the DV specificity of PAX dual activity is dictated by its cellular context, particularly BMP exposure. Ultimately, there is a dorsoventral specificity in the modulation of PAX dual activity, dictated by the dependence of PAX-mediated activation on the cellular context established by BMP signalling. Since BMP signalling is spatially restricted to dorsal regions, PAX activation is likewise confined, thereby providing a spatial input to the regulation of its target genes.

The generic activity of PAX3/7 across their expression domain is transcriptional repression, echoing the critical role of repression in TF-mediated spinal cord patterning (Sagner and Briscoe 2019). However, this repression does not universally target CRMs near genes regulating all alternative fates to PAX-expressing cells, as observed for ventral TFs (Kutejova et al. 2016; Nishi et al. 2015). For example, genes specifying SHH-dependent ventral identities (p2, p3, pMN) remained unaffected by the PAX3 and PAX7 modulation (Table S1). Instead, PAX3/7-bound silencers are primarily associated with p0 and p1 progenitor regulators and their V0/V1 neuronal derivatives. Furthermore, PAX3 is transiently expressed in p0 and p1 progenitors (Moore et al. 2013), suggesting that repression may be lifted during the differentiation, potentially influencing its timing.

The specification of dp4–dp6 and p0–p1 identities has long been debated, as classical neural tube signalling pathways, such as Shh and BMPs, do not fully explain their establishment (Pierani et al. 2001; Moore et al. 2013; Zagorski et al. 2017; Sasai et al. 2014; Pierani et al. 1999). These intermediate spinal cord regions are only transiently exposed to noisy dorsal (BMP) and ventral (Shh) morphogen gradients (Zagorski et al. 2017). Moreover, exposure to these signals often inhibits intermediate fate specification, suggesting these identities emerge as default states of spinal progenitors (Pierani et al. 1999; Zagorski et al. 2017; Sasai et al. 2014). Consistently, in our organoid differentiation experiments, dp4–dp6 and p0–p1 identities arise in the absence of factors modulating BMP or Shh signalling ((Duval et al. 2019) and this study). Our findings reveal that PAX-mediated repression imposes a choice between these identities. In the absence of PAX factors, cell fate specification shifts toward p0–p1 states (Gard et al. 2017; Mansouri and Gruss 1998) and this study), even in the presence of BMPs. Among PAX-bound silencers, several are located near *Dbx1* and *Prdm12*, key determinants of these ventral identities, whose deregulation in the absence of PAX could explain the ventralisation of dorsal identities (Thélie et al. 2015; Pierani et al. 2001).

PAX3/7-mediated transcriptional repression is particularly striking, as PAX proteins are predominantly recognized for their activator role in cellular differentiation across embryonic tissues (Mayran et al. 2015). However, our findings may extend to these systems, as studies on a limited number of CRMs and co-repressors have suggested a repressive function for PAX proteins (Choi et al. 2014; Agarwal et al. 2022). In the spinal cord, our results suggest that this repression involves modulating H3K27me3 deposition, a process typically governed by the Polycomb complexes. PAX2 is known to recruit these complexes through Groucho/TLE co-repressors, binding through a conserved octapeptide domain shared with PAX3 and PAX7 (Eberhard et al. 2000; Patel et al. 2012). Several lines of evidence support this mechanism in PAX3- and PAX7-mediated repression: i) PAX7 biochemically interacts with TLE4 (Muhr et al. 2001). ii) Deleting the octapeptide domain enhances PAX3 activator potential in myoblasts (Magli et al. 2013). iii) Inhibiting TLE activity in the neural tube of chicken embryos leads to ectopic expression of genes, such as *Dbx1*, normally repressed by PAX (Muhr et al. 2001). Moreover, in light of a study on the Dorsal protein in *Drosophila*, we propose that such a PAX-TLE repression mechanism may also enable simultaneous activation and repression in dp1– dp3 cells (Ratnaparkhi et al. 2006). Like Dorsal, PAX octapeptide diverges from the canonical high-affinity Groucho/TLE-binding sequence, likely reducing its affinity for TLE (Eberhard et al. 2000; Muhr et al. 2001). This lower affinity may make PAX-TLE interactions highly context-dependent, fine-tuning PAX-mediated transcriptional regulation (Ratnaparkhi et al. 2006).

In contrast to PAX-mediated repression, the transcriptional activation mechanisms of PAX proteins in dp1 and dp3 progenitors align closely with previous findings (Kawabe et al. 2012; Diao et al. 2012; McKinnell et al. 2008; Budry et al. 2012; Magli et al. 2019; Gouhier et al. 2024). In the spinal cord, PAX proteins are required for opening or maintaining chromatin accessibility at CRMs and, in some cases, for depositing the enhancer mark H3K4me2. This mirrors studies on PAX7 recruitment to CRMs, where it functions as a pioneer factor, recruiting an H3K9me2 demethylase and the MLL methyltransferase complex to modify H3K4 (Gouhier et al. 2024). Following cell division, which enables the dissociation of PAX3/7-bound CRMs from the nuclear lamina, PAX7 can recruit SWI-SNF nucleosome remodelers and the coactivator P300.

The restriction of PAX3- and PAX7-mediated transcriptional activation to BMP-exposed progenitors is intriguing. While the exact mechanisms remain to be uncovered, several promising hypotheses emerge from our study. A direct interaction between PAX proteins and SMADs, the BMP transcriptional effectors, seems unlikely, as PAX3/7 activity persists for up to four days—far exceeding the transient BMP peak, which declines within an hour (Duval et al. 2019). Moreover, PAX3/7-bound CRMs show no enrichment for SMAD-binding motifs. The presence of PAX3/7-bound CRMs specific to dp1-dp3 progenitors suggests that indirect BMP effectors may facilitate PAX recruitment. Notably, the enrichment of ZIC motifs in these CRMs suggests a plausible BMP-specific PAX-ZIC partnership, further supported by the exclusive expression of ZIC1, ZIC2, and ZIC5 in BMP-exposed progenitors (Inoue et al. 2004), and the demonstrated interaction between PAX3 and ZIC2 in regulating a *Myf5* CRM (Himeda et al. 2013). Alternatively, BMP effectors may modulate PAX activity on already bound CRMs rather than influencing its recruitment. Supporting this, nearly half of PAX-activated CRMs in BMP-treated conditions were PAX3/7-bound regardless of BMP exposure and the *P34::tk::LacZ* reporter transgene, containing only PAX-binding motifs, is restricted dorsally. Post-translational modifications (e.g., acetylation (Ichi et al. 2011), SUMOylation (Luan et al. 2012), phosphorylation (Iyengar et al. 2014)) or alternative splicing could regulate PAX activity (Mayran et al. 2015; Barber et al. 1999; Bascunana et al. 2023), and so could the recruitment of specific cofactors such as ZIC1 (Himeda et al. 2013) or Lef/TCF (Werner et al. 2007).

## Materials and Methods

### Mouse lines

Mice carrying the *Pax3* knock-in *GFP* null allele (*PAX3^GFP^*), conditional PAX3-FOXO1 (*PAX3^PAX3-FOXO1^)*, Pax3-EnR (*PAX3^Pax3-EnR^)*, and *Pax7* knock-in *LacZ* null (*PAX7^LacZ^*) alleles and the *P34::tk::LacZ* reporter transgene have been previously described (Bajard et al. 2006; Mansouri et al. 1996; Relaix et al. 2003; Zalc et al. 2015). The *PAX3-FOXO1* and *Pax3-EnR* alleles were expressed from the *Pax3* locus upon activation of Cre recombinase, driven by the zygote-specific *PGK* enhancer (Lallemand et al. 1998). All experiments were conducted according to the Institutional Animal Care and Use Committee of the university Paris Cité.

### mESC cell lines maintenance and differentiation in organoids

All mouse ESC lines were maintained on mitotically inactive primary mouse embryo fibroblasts in D-MEM Glutamax supplemented with 10% ESC-qualified foetal bovine serum (Millipore), L-glutamine, non-essential amino acid, nucleosides, 0.1 mM β-mercaptoethanol (Life technologies) and 1000 U.ml^−1^ leukemia inhibitory factor (Millipore). To set mESC embryoid bodies differentiation, cells were trypsinised and placed twice onto gelatinised tissue culture plates to remove feeders. From 5×10^5^ cells.ml^-1^ to 1×10^6^ cells.ml^-1^ were placed in ultra-low attachment petri dishes (Corning) and in Advanced Dulbecco’s Modified Eagle/ F12 and Neurobasal media (1:1, Life technologies) supplemented with 1X B27 devoid of Vitamin A and 1X N2 (Life technologies), 2 mM L-glutamine (Life technologies), 0.1 mM β-mercaptoethanol, 100U/mL penicillin and 100U/mL streptomycin (Life technologies). At this concentration of cells, small EB were formed from day 1 of differentiation, and grown as such up to day 7 of differentiation; medium was changed every day from day 2 onwards. Chemical drugs to inhibit or activate key developmental important signalling pathways were used at the following concentrations: 3 μM CHIR99021 (Tocris or Axon Medchem), 10 nM Retinoic Acid (Sigma), 5 ng.ml^-1^ BMP4 (R&D).

### Generation of transgenic mESC

The *P34::tk::LacZ* mESC line was derived from mouse blastocysts following a previously published protocol (Kanda et al. 2012). To create *Pax3^-/-^*, *Pax7^-/-^*, or *Pax3^-/-^; Pax7^-/-^* lines, we employed CRISPR-Cas9-mediated deletion of the first two exons and half of the third exon for each gene (this deletion eliminates five ATG codons and creates a frameshift). The guide RNAs used are provided in Table S4. For *Pax3^-/-^*, following passage, 1×10^6^ cells were plated and were co-transfected with *pX459* plasmid enabling transient expression of *Cas9* and the sgRNA for *Pax3* first exon, as well as another *pX459* plasmid expressing the sgRNA for *Pax3* third exon The plasmid *pSpCas9(BB)-2A-Puro (PX459)* V2.0 was a gift from Feng Zhang (Addgene plasmid # 62988; http://n2t.net/addgene:62988; RRID: Addgene_62988). Transfection was performed using Lipofectamine 2000 (Life technologies, Carlsbad, CA, USA) according to the manufacturer’s instructions. Twenty-four hours later, puromycin (2 μg/ml) was added to the medium. After two days, the cells were diluted 1:100 and replated. Four days later, 96 isolated colonies were picked and split into two portions. After removal of feeders, DNA was extracted using the RedExtract-N-Amp kit, and the clones were genotyped via PCR (Table S5). Three independent homozygous clones per genotype were selected. The same strategy was applied for *Pax7^-/-^*. For *Pax3^-/-^; Pax7^-/-^* lines, cells were co-transfected with the four plasmids. To generate *Flag-Pax3* cell line, cells were co-transfected with the basal *pX459* and a single-stranded donor DNA enabling the integration of *3xFLAG* tag upstream of the first ATG codon of *Pax3*, using the same transfection protocol (Table S4). After genotyping, only one clone exhibited the correct integration of the tag; this clone was amplified and used for CUT&Tag experiments. The same approach was used to generate the *Flag-Pax7* cell line.

### Reporter constructs and in ovo electroporation

Enhancers or silencers bound by PAX3/7 were amplified by PCR and inserted upstream of the minimal *adenovirus Major Late Promoter* (MLP) and *H2B-Turquoise* for enhancers, or upstream of the *EF1alpha* constitutive promoter and *H2B-Turquoise* for silencers, using the Takara In-Fusion kit. The mutant versions of CRMs were obtained by PCR-mediated directed mutagenesis using the QuickChange Multi Site Directed Mutagenesis Kit from Agilent. Reporter plasmids (1 μg/μl) and a *pCAG* plasmid enabling constitutive expression of the mCherry reporter (0.3 μg/μl) were electroporated into Hamburger and Hamilton (HH) stage 10–11 chick embryos following established protocols (Briscoe et al. 2000). Embryos were dissected at the indicated stage in cold 1X PBS.

### Immunohistochemistry on histological sections

Mouse and chick embryos were fixed in 4% paraformaldehyde for 45 minutes to 2 hours at 4°C. Fixed embryos were cryoprotected by equilibration in 15% sucrose, embedded in gelatine, cryosectioned at 14 µm, and processed for immunostaining (Briscoe et al. 2000). The fixation, embedding, and cryosectioning of EBs were conducted as previously described (Maury et al. 2015), and immunolabelling was performed using the same protocol as for embryo sections. Details of the antibodies are provided in Table S6. Images were carried out using either a Leica TCS SP5 confocal microscope or a Zeiss LSM 980, and images were processed with Adobe Photoshop 7.0 (Adobe Systems, San Jose, CA, USA) or ImageJ v.1.43g (NIH). All scale bars represent 50 µm. In mouse embryos, quantifications were performed at brachial levels on usually more than 3 embryos and on 2 to 4 transverse sections per embryo.

### Quantification based on immunostaining signals

The number of cells per section of mouse spinal cords was determined using the Cell Counter plugin in ImageJ v.1.43g (NIH). The fluorescence intensity of β-galactosidase, GFP, Turquoise, and mCherry in chick embryos, as well as the evaluation of dorsal–ventral boundary positions, were determined as described in (Balaskas et al. 2012). The number of cells immuno-labelled in organoids was calculated using Cell Profiler (Broad Institute) after nuclei segmentation based on the DAPI fluorescence signal and a defined signal intensity threshold, expressed as a percentage of all detected nuclei. Fluorescence signal intensity levels of PAX3, PAX7, or phosphorylated SMAD1/5/9 per cell were evaluated using CellProfiler, followed by background subtraction. Background values were determined in EBs not treated with BMP4 for phosphorylated SMAD1/5/9 or in EBs generated from *Pax3^-/-^; Pax7^-/-^*cell lines. Statistical analyses and graphs were performed using GraphPad Prism software. For each graph, the biological replicates (embryos or ESC clones, typically ≥ 3) and technical replicates (individual sections) were displayed. Non-parametric t-tests (Mann–Whitney U test) were used for pairwise comparisons between conditions. P-values are denoted as follows: * for p ≤ 0.05, ** for p ≤ 0.01, *** for p ≤ 0.001, and **** for p ≤ 0.0001. All quantified data can be found in Table S7.

### RNA extraction, qPCR and sequencing

RNAs were extracted using the NucleoSpin RNA kit (Macherey-Nagel) following manufacturer’s instructions, elution was done with RNase-free water and concentrations were measured with DeNovixDS-11-FX series. For real time quantitative PCR, cDNA were synthesised using SuperScript IV (Thermo Fischer Scientific), random primers and oligo dT, then SYBR Green I Master (Roche) and the LightCycler 480 II (Roche) were used. PCR primers were designed using Primer-Blast (Table S8). Levels of expression per gene for a given time point was measured in biological duplicates or triplicates. Gene expression levels were normalised to the levels of *Ywhaz* mRNA in ESCs. For transcriptomic analysis, RNAs were sent to BGI (Hong-Kong) where libraries were prepared and their quality checked using a Bioanalyzer Agilent 2100. 0.2-0.5µg was then either sequenced on a DBN sequencing system on paired-end reads 150 base runs (D6 organoids). After demultiplexing, reads were trimmed and mapped on the mm10 mouse genome (BGI).

### Transcriptomic analyses

Pairwise differential analysis of RNA-seq data was performed using Deseq2 (Love et al. 2014) (BGI). Differentially expressed genes were defined as those with a mean expression >5 FPKM in at least one group, an absolute fold change > 1.5, and a q-value <0.05 after Benjamini-Hochberg correction for multiple-testing were considered to be significant. Volcano plotS were generated using ggplot2 (Wickham 2016). Line plots were generated in GraphPad Prism. Ad-hoc enrichment analyses between differentially expressed genes, and gene sets defining DV populations (Table S1) were performed using hypergeometric tests (phyper function on R). The lists of genes coding for DV markers genes (Table S1) were generated based on literature reviewing and were used to project the effects of the PAXs onto the acquisition of DV identities. Heatmaps of relative expression levels were generated using the plotHeatmap function in R.

### ATAC-Seq, CUT&Tag and sequencing

ATAC-Seq was performed as in (Buenrostro et al. 2013). We used between 50 000 cells per condition, and a homemade purified pA-TN5 protein. CUT&Tag was performed as in (Kaya-Okur et al. 2020), following the procedure described in “Bench top CUT&Tag V.2” available on protocols.io. We used 500 000 cells per sample and the anti-FLAG (Sigma-Aldrich F3165-5MG; 1/50) for CUT&Tag on transcription factors, or 200 000 nuclei that were previously fixed and frozen following the procedure described in (Kaya-Okur et al. 2020) with anti-H3K4me2 (Abcam ab176878; 1/100), anti-H3K27me3 (Cell Signalling 9733S; 1/100) primary antibodies; and anti-rabbit IgG (ABIN6923140) or anti-mouse IgG (ABIN6923141) secondary antibodies. We used a homemade purified pA-TN5 protein. Libraries were analysed using 4200 TapeStation system (Agilent) and sequenced 100pb (paired-end) with DNBSEQ-G400 (BGI). These data can be found in Table S2.

### Bioinformatic analyses

The quality of paired-end read sequences was assessed using FastQC (v0.11.9). Low quality nucleotides and Illumina adapter sequences were removed using Trimmomatic (v0.39)(Bolger et al. 2014) and PCR duplicate reads were removed using BBmap clumpify (v38.87)(Bushnell 2014). Filtered reads were aligned to the mm10 reference genome using Bowtie2 (v2.4.5)(Langmead and Salzberg 2012) and parameters “--local --very-sensitive --no-mixed --dovetail --no-discordant --phred33 -I 10 -X 700”. Genome-wide coverage tracks (bigwigs) were generated using deeptools (v3.5.1)(Ramírez et al. 2016) bamCoverage and parameter “--normalizeUsing CPM --blackListFileName blacklisted_regions.fa –ignoreForNormalization chrX chrM chrY --binsize 1”, removing blacklisted regions defined by the Kundaje lab. Peaks were called using MACS2 (v 2.2.7.1)(Zhang et al. 2008) with parameters: -f BAMPE -broad -g 1.87e9 and manually curated. Reads coverage on defined CRMs was calculated using VisRseq (v0.9.2) by taking the signal on each CRM, then normalised by library size. Differential analysis on PAX bound peaks was performed using Deseq2 (v1.42.1)(Love et al. 2014), allowing the identification of three CRMs categories (abs(fold change) > 1.5; adjusted p-value < 0.05). For ATAC-Seq, H3K4me2 and H3K27me3, peaks were called in every condition using the same MACS2 parameters, and for H3K27me3, peaks within 5000bp of each other were merged. Peaks obtained for ATAC in the different conditions were merged to form a universal of list of peaks, and the same was done for the other chromatin parameters. Subsequently, enrichment was calculated using VisRseq, and differential analyses were performed using DeSeq2, allowing us to define differentially open, activated or repressed regions. Using bedtools intersect between PAX CRM and these marks data, we could identify the chromatin state of the different PAX CRMs. Heatmaps and average plots were generated using deeptools (v3.5.1)(Ramírez et al. 2016) computeMatrix followed by plotHeatmap or plotProfile. PCA plots were generated with plotPCA command in R. The association between PAX3/7-bound CRM and gene was performed using GREAT (McLean et al. 2010). Heatmaps of relative expression levels were generated using the plotHeatmap function in R. UCSC Genome Browser (Kent et al. 2002) track data hubs (Raney et al. 2014) were used to display data over loci of interest.

### Motifs analyses

Position weight matrices (PWMs) for TFs binding sites were taken from the JASPAR 2022 database (Castro-Mondragon et al. 2022), to which we added PWM created *de novo* from PAX3, PAX7 or PAX-FOXO1 ChIP-Sequencing data previously published (Manceau et al. 2022) (Manceau et al. 2022). Relative enrichment of those PWM was calculated in PAX3/7 CUT&Tag data using AME (McLeay and Bailey 2010) from MEME-suite. The logos of the PAX3/7 binding motifs were generated from PWM using ggseqlogo (Wagih 2017).

### Resource availability

All the raw and processed data associated with this study will be submitted to the NCBI Gene Expression Omnibus (GEO) (GSE288918).

## Acknowledgments

PGH is a CNRS research director; VR and FR are INSERM research directors; BD is an INSERM engineer. RR received a three-year PhD fellowship from the University of Paris, and his fourth year of PhD was supported by the Labex Who Am I? (ANR-11-LABX-0071). This work was funded by grants to VR from the CNRS/INSERM ATIP-AVENIR program, the Ligue Nationale Contre le Cancer (PREAC2020.LCC/MC; PREAC2016.LCC; RS20/75-114), and the Agence Nationale de la Recherche (ANR) grant AetioSpinoid (ANR-23-CE16-0026-01), as well as by funding to FR from the Labex REVIVE (ANR-10-LABX-73). We sincerely thank C. Birchmeier, T. Müller, and S. Garel for providing antibodies. We also acknowledge the ImagoSeine core facility at the Institut Jacques Monod, a member of France-BioImaging (ANR-10-INBS-04) and certified by GIS-IBiSA. We are grateful to the biological service staff of the CDTA and Buffon animal housing for help with the mouse colonies. Finally, we are grateful to S. Nedelec and N. Konstantinides for his critical comments on the manuscript.

## Author contributions

**Conceptualization:** R.R., P.G.H., V.R., C.D.D.

**Experiments:** PAX modified ESC lines: P.G.H., R.R.; CUT&Tags: P.G.H., R.R.; CRM activity in chick embryos: P.G.H., R.R.; organoid phenotypes: R.R., P.G.H., T.H., C.D.D., V.R., Mouse embryos phenotypes: V.R., G.G.C.; *P34::tk::nLacZ* ESC derivation: B.D.L., V.R. in F.R.’s lab.

**Bioinformatic analyses:** R.R., J.R.A.

**Writing:** primary draft V.R., R.R., P.G.H.;

**Revision:** C.D.D., T.H.

**Supervision:** P.G.H., V.R.

**Funding acquisition:** V.R., F.R.

## References

Agarwal M, Bharadwaj A, Mathew SJ. 2022. TLE4 regulates muscle stem cell quiescence and skeletal muscle differentiation. Journal of Cell Science 135: jcs256008.

Alvarez-Medina R, Cayuso J, Okubo T, Takada S, Martí E. 2008. Wnt canonical pathway restricts graded Shh/Gli patterning activity through the regulation of Gli3 expression. Development 135: 237–247.

Andrews MG, del Castillo LM, Ochoa-Bolton E, Yamauchi K, Smogorzewski J, Butler SJ. 2017. BMPs direct sensory interneuron identity in the developing spinal cord using signal-specific not morphogenic activities. eLife 6: e30647.

Bailey PJ, Klos JM, Andersson E, Karlén M, Källström M, Ponjavic J, Muhr J, Lenhard B, Sandelin A, Ericson J. 2006. A global genomic transcriptional code associated with CNS-expressed genes. Experimental Cell Research 312: 3108–3119.

Bajard L, Relaix F, Lagha M, Rocancourt D, Daubas P, Buckingham ME. 2006. A novel genetic hierarchy functions during hypaxial myogenesis: Pax3 directly activates *Myf5* in muscle progenitor cells in the limb. Genes Dev 20: 2450–2464.

Balaskas N, Ribeiro A, Panovska J, Dessaud E, Sasai N, Page KM, Briscoe J, Ribes V. 2012. Gene Regulatory Logic for Reading the Sonic Hedgehog Signaling Gradient in the Vertebrate Neural Tube. Cell 148: 273–284.

Barber TD, Barber MC, Cloutier TE, Friedman TB. 1999. *PAX3* gene structure, alternative splicing and evolution. Gene 237: 311–319.

Bascunana V, Pelletier A, Gouhier A, Bemmo A, Balsalobre A, Drouin J. 2023. Chromatin opening ability of pioneer factor Pax7 depends on unique isoform and C-terminal domain. Nucleic Acids Res 51: 7254–7268.

Bolger AM, Lohse M, Usadel B. 2014. Trimmomatic: a flexible trimmer for Illumina sequence data. Bioinformatics 30: 2114–2120.

Borromeo MD, Meredith DM, Castro DS, Chang JC, Tung K-C, Guillemot F, Johnson JE. 2014. A transcription factor network specifying inhibitory versus excitatory neurons in the dorsal spinal cord. Development 141: 2803–2812.

Briscoe J, Pierani A, Jessell TM, Ericson J. 2000. A Homeodomain Protein Code Specifies Progenitor Cell Identity and Neuronal Fate in the Ventral Neural Tube. Cell 101: 435–445.

Budry L, Balsalobre A, Gauthier Y, Khetchoumian K, L’Honore A, Vallette S, Brue T, Figarella-Branger D, Meij B, Drouin J. 2012. The selector gene Pax7 dictates alternate pituitary cell fates through its pioneer action on chromatin remodeling. Genes & Development 26: 2299–2310.

Buenrostro JD, Giresi PG, Zaba LC, Chang HY, Greenleaf WJ. 2013. Transposition of native chromatin for fast and sensitive epigenomic profiling of open chromatin, DNA-binding proteins and nucleosome position. Nat Methods 10: 1213–1218.

Bushnell B. 2014. BBMap: A Fast, Accurate, Splice-Aware Aligner. https://escholarship.org/uc/item/1h3515gn (Accessed May 6, 2024).

Castro-Mondragon JA, Riudavets-Puig R, Rauluseviciute I, Berhanu Lemma R, Turchi L, Blanc-Mathieu R, Lucas J, Boddie P, Khan A, Manosalva Pérez N, et al. 2022. JASPAR 2022: the 9th release of the open-access database of transcription factor binding profiles. Nucleic Acids Research 50: D165–D173.

Chang JC, Meredith DM, Mayer PR, Borromeo MD, Lai HC, Ou Y-H, Johnson JE. 2013. Prdm13 Mediates the Balance of Inhibitory and Excitatory Neurons in Somatosensory Circuits. Developmental Cell 25: 182–195.

Choi M, Ryu S, Hao R, Wang B, Kapur M, Fan C, Yao T. 2014. HDAC 4 promotes P ax7-dependent satellite cell activation and muscle regeneration. EMBO Reports 15: 1175–1183.

Delile J, Rayon T, Melchionda M, Edwards A, Briscoe J, Sagner A. 2019. Single cell transcriptomics reveals spatial and temporal dynamics of gene expression in the developing mouse spinal cord. Development dev.173807.

Diao Y, Guo X, Li Y, Sun K, Lu L, Jiang L, Fu X, Zhu H, Sun H, Wang H, et al. 2012. Pax3/7BP Is a Pax7- and Pax3-Binding Protein that Regulates the Proliferation of Muscle Precursor Cells by an Epigenetic Mechanism. Cell Stem Cell 11: 231–241.

Duval N, Vaslin C, Barata T, Frarma Y, Contremoulins V, Baudin X, Nédélec S, Ribes V. 2019. BMP4 patterns Smad activity and generates stereotyped cell fate organisation in spinal organoids. Development dev.175430.

Eberhard D, Jiménez G, Heavey B, Busslinger M. 2000. Transcriptional repression by Pax5 (BSAP) through interaction with corepressors of the Groucho family. EMBO J 19: 2292– 2303.

Ernst J, Kheradpour P, Mikkelsen TS, Shoresh N, Ward LD, Epstein CB, Zhang X, Wang L, Issner R, Coyne M, et al. 2011. Mapping and analysis of chromatin state dynamics in nine human cell types. Nature 473: 43–49.

Gard C, Gonzalez Curto G, Frarma YE-M, Chollet E, Duval N, Auzié V, Auradé F, Vigier L, Relaix F, Pierani A, et al. 2017. Pax3- and Pax7-mediated Dbx1 regulation orchestrates the patterning of intermediate spinal interneurons. Developmental Biology 432: 24–33.

Gouhier A, Dumoulin-Gagnon J, Lapointe-Roberge V, Harris J, Balsalobre A, Drouin J. 2024. Pioneer factor Pax7 initiates two-step cell-cycle-dependent chromatin opening. Nat Struct Mol Biol 31: 92–101.

Goulding MD, Chalepakis G, Deutsch U, Erselius JR, Gruss P. 1991. Pax-3, a novel murine DNA binding protein expressed during early neurogenesis. The EMBO Journal 10: 1135–1147.

Himeda CL, Barro MV, Emerson CP. 2013. Pax3 synergizes with Gli2 and Zic1 in transactivating the Myf5 epaxial somite enhancer. Developmental Biology 383: 7–14.

Ichi S, Boshnjaku V, Shen Y-W, Mania-Farnell B, Ahlgren S, Sapru S, Mansukhani N, McLone DG, Tomita T, Mayanil CSK. 2011. Role of Pax3 acetylation in the regulation of Hes1 and Neurog2 ed. M. Bronner-Fraser. MBoC 22: 503–512.

Inoue T, Hatayama M, Tohmonda T, Itohara S, Aruga J, Mikoshiba K. 2004. Mouse Zic5 deficiency results in neural tube defects and hypoplasia of cephalic neural crest derivatives. Developmental Biology 270: 146–162.

Iyengar AS, Miller PJ, Loupe JM, Hollenbach AD. 2014. Phosphorylation of PAX3 contributes to melanoma phenotypes by affecting proliferation, invasion, and transformation. Pigment Cell Melanoma Res 27: 846–848.

Kanda A, Sotomaru Y, Shiozawa S, Hiyama E. 2012. Establishment of ES Cells from Inbred Strain Mice by Dual Inhibition (2i). J Reprod Dev 58: 77–83.

Katsuda T, Sussman JH, Zaret KS, Stanger BZ. 2024. The yin and yang of pioneer transcription factors: Dual roles in repression and activation. BioEssays 46: 2400138.

Kawabe Y, Wang YX, McKinnell IW, Bedford MT, Rudnicki MA. 2012. Carm1 Regulates Pax7 Transcriptional Activity through MLL1/2 Recruitment during Asymmetric Satellite Stem Cell Divisions. Cell Stem Cell 11: 333–345.

Kaya-Okur HS, Janssens DH, Henikoff JG, Ahmad K, Henikoff S. 2020. Efficient low-cost chromatin profiling with CUT&Tag. Nat Protoc 15: 3264–3283.

Kent WJ, Sugnet CW, Furey TS, Roskin KM, Pringle TH, Zahler AM, Haussler and D. 2002. The Human Genome Browser at UCSC. Genome Res 12: 996–1006.

Kicheva A, Bollenbach T, Ribeiro A, Valle HP, Lovell-Badge R, Episkopou V, Briscoe J. 2014. Coordination of progenitor specification and growth in mouse and chick spinal cord. Science 345: 1254927.

Koromila T, Stathopoulos A. 2017. Broadly expressed repressors integrate patterning across orthogonal axes in embryos. Proc Natl Acad Sci USA 114: 8295–8300.

Kutejova E, Sasai N, Shah A, Gouti M, Briscoe J. 2016. Neural Progenitors Adopt Specific Identities by Directly Repressing All Alternative Progenitor Transcriptional Programs. Developmental Cell 36: 639–653.

Lai HC, Seal RP, Johnson JE. 2016. Making sense out of spinal cord somatosensory development. Development 143: 3434–3448.

Lallemand Y, Luria V, Haffner-Krausz R, Lonai P. 1998. Maternally expressed PGK-Cre transgene as a tool for early and uniform activation of the Cre site-specific recombinase. Transgenic Res 7: 105–112.

Langmead B, Salzberg SL. 2012. Fast gapped-read alignment with Bowtie 2. Nat Methods 9: 357–359.

Le Dréau G, Garcia-Campmany L, Rabadán MA, Ferronha T, Tozer S, Briscoe J, Martí E. 2012. Canonical BMP7 activity is required for the generation of discrete neuronal populations in the dorsal spinal cord. Development 139: 259–268.

Lodato MA, Ng CW, Wamstad JA, Cheng AW, Thai KK, Fraenkel E, Jaenisch R, Boyer LA. 2013. SOX2 Co-Occupies Distal Enhancer Elements with Distinct POU Factors in ESCs and NPCs to Specify Cell State. PLoS Genet 9: e1003288.

Love MI, Huber W, Anders S. 2014. Moderated estimation of fold change and dispersion for RNA-seq data with DESeq2. Genome Biology 15: 550.

Luan Z, Liu Y, Stuhlmiller TJ, Marquez J, García-Castro MI. 2012. SUMOylation of Pax7 is essential for neural crest and muscle development. Cell Mol Life Sci 70: 1793–1806.

Lukoseviciute M, Gavriouchkina D, Williams RM, Hochgreb-Hagele T, Senanayake U, Chong-Morrison V, Thongjuea S, Repapi E, Mead A, Sauka-Spengler T. 2018. From Pioneer to Repressor: Bimodal foxd3 Activity Dynamically Remodels Neural Crest Regulatory Landscape In Vivo. Developmental Cell 47: 608–628.e6.

Magli A, Baik J, Pota P, Cordero CO, Kwak I-Y, Garry DJ, Love PE, Dynlacht BD, Perlingeiro RCR. 2019. Pax3 cooperates with Ldb1 to direct local chromosome architecture during myogenic lineage specification. Nat Commun 10: 2316.

Magli A, Schnettler E, Rinaldi F, Bremer P, Perlingeiro RCR. 2013. Functional Dissection of Pax3 in Paraxial Mesoderm Development and Myogenesis. STEM CELLS 31: 59–70.

Manceau L, Richard Albert J, Lollini P-L, Greenberg MVC, Gilardi-Hebenstreit P, Ribes V. 2022. Divergent transcriptional and transforming properties of PAX3-FOXO1 and PAX7-FOXO1 paralogs. PLoS Genet 18: e1009782.

Mansouri A, Gruss P. 1998. Pax3 and Pax7 are expressed in commissural neurons and restrict ventral neuronal identity in the spinal cord. Mechanisms of Development 78: 171–178.

Mansouri A, Stoykova A, Torres M, Gruss P. 1996. Dysgenesis of cephalic neural crest derivatives in *Pax7* − */* − mutant mice. Development 122: 831–838.

Maury Y, Côme J, Piskorowski RA, Salah-Mohellibi N, Chevaleyre V, Peschanski M, Martinat C, Nedelec S. 2015. Combinatorial analysis of developmental cues efficiently converts human pluripotent stem cells into multiple neuronal subtypes. Nat Biotechnol 33: 89–96.

Mayran A, Khetchoumian K, Hariri F, Pastinen T, Gauthier Y, Balsalobre A, Drouin J. 2018. Pioneer factor Pax7 deploys a stable enhancer repertoire for specification of cell fate. Nat Genet 50: 259–269.

Mayran A, Pelletier A, Drouin J. 2015. Pax factors in transcription and epigenetic remodelling. Seminars in Cell & Developmental Biology 44: 135–144.

Mayran A, Sochodolsky K, Khetchoumian K, Harris J, Gauthier Y, Bemmo A, Balsalobre A, Drouin J. 2019. Pioneer and nonpioneer factor cooperation drives lineage specific chromatin opening. Nat Commun 10: 3807.

McKinnell IW, Ishibashi J, Le Grand F, Punch VGJ, Addicks GC, Greenblatt JF, Dilworth FJ, Rudnicki MA. 2008. Pax7 activates myogenic genes by recruitment of a histone methyltransferase complex. Nat Cell Biol 10: 77–84.

McLean CY, Bristor D, Hiller M, Clarke SL, Schaar BT, Lowe CB, Wenger AM, Bejerano G. 2010. GREAT improves functional interpretation of cis-regulatory regions. Nat Biotechnol 28: 495–501.

McLeay RC, Bailey TL. 2010. Motif Enrichment Analysis: a unified framework and an evaluation on ChIP data. BMC Bioinformatics 11: 165.

Moore S, Ribes V, Terriente J, Wilkinson D, Relaix F, Briscoe J. 2013. Distinct Regulatory Mechanisms Act to Establish and Maintain Pax3 Expression in the Developing Neural Tube ed. C. Desplan. PLoS Genet 9: e1003811.

Muhr J, Andersson E, Persson M, Jessell TM, Ericson J. 2001. Groucho-Mediated Transcriptional Repression Establishes Progenitor Cell Pattern and Neuronal Fate in the Ventral Neural Tube. Cell 104: 861–873.

Nakazaki H, Reddy AC, Mania-Farnell BL, Shen Y-W, Ichi S, McCabe C, George D, McLone DG, Tomita T, Mayanil CSK. 2008. Key basic helix–loop–helix transcription factor genes Hes1 and Ngn2 are regulated by Pax3 during mouse embryonic development. Developmental Biology 316: 510–523.

Nishi Y, Zhang X, Jeong J, Peterson KA, Vedenko A, Bulyk ML, Hide WA, McMahon AP. 2015. A direct fate exclusion mechanism by Sonic hedgehog-regulated transcriptional repressors. Development dev.124636.

Oosterveen T, Kurdija S, Alekseenko Z, Uhde CW, Bergsland M, Sandberg M, Andersson E, Dias JM, Muhr J, Ericson J. 2012. Mechanistic Differences in the Transcriptional Interpretation of Local and Long-Range Shh Morphogen Signaling. Developmental Cell 23: 1006–1019.

Oosterveen T, Kurdija S, Enstero M, Uhde CW, Bergsland M, Sandberg M, Sandberg R, Muhr J, Ericson J. 2013. SoxB1-driven transcriptional network underlies neural-specific interpretation of morphogen signals. Proceedings of the National Academy of Sciences 110: 7330–7335.

Patel SR, Bhumbra SS, Paknikar RS, Dressler GR. 2012. Epigenetic Mechanisms of Groucho/Grg/TLE Mediated Transcriptional Repression. Molecular Cell 45: 185–195.

Pierani A, Brenner-Morton S, Chiang C, Jessell TM. 1999. A Sonic Hedgehog–Independent, Retinoid-Activated Pathway of Neurogenesis in the Ventral Spinal Cord. Cell 97: 903–915.

Pierani A, Moran-Rivard L, Sunshine MJ, Littman DR, Goulding M, Jessell TM. 2001. Control of Interneuron Fate in the Developing Spinal Cord by the Progenitor Homeodomain Protein Dbx1. Neuron 29: 367–384.

Ramírez F, Ryan DP, Grüning B, Bhardwaj V, Kilpert F, Richter AS, Heyne S, Dündar F, Manke T. 2016. deepTools2: a next generation web server for deep-sequencing data analysis. Nucleic Acids Res 44: W160–165.

Ramos R, Swedlund B, Ganesan AK, Morsut L, Maini PK, Monuki ES, Lander AD, Chuong C-M, Plikus MV. 2024. Parsing patterns: Emerging roles of tissue self-organization in health and disease. Cell 187: 3165–3186.

Raney BJ, Dreszer TR, Barber GP, Clawson H, Fujita PA, Wang T, Nguyen N, Paten B, Zweig AS, Karolchik D, et al. 2014. Track data hubs enable visualization of user-defined genome-wide annotations on the UCSC Genome Browser. Bioinformatics 30: 1003–1005.

Ratnaparkhi GS, Jia S, Courey AJ. 2006. Uncoupling Dorsal-mediated activation from Dorsal-mediated repression in the *Drosophila* embryo. Development 133: 4409–4414.

Rekler D, Ofek S, Kagan S, Friedlander G, Kalcheim C. 2024. Retinoic acid, an essential component of the roof plate organizer, promotes the spatiotemporal segregation of dorsal neural fates. Development 151: dev202973.

Relaix F, Polimeni M, Rocancourt D, Ponzetto C, Schäfer BW, Buckingham M. 2003. The transcriptional activator PAX3-FKHR rescues the defects of *Pax3* mutant mice but induces a myogenic gain-of-function phenotype with ligand-independent activation of Met signaling in vivo. Genes Dev 17: 2950–2965.

Sagner A, Briscoe J. 2019. Establishing neuronal diversity in the spinal cord: a time and a place. Development 146: dev182154.

Sasai N, Kutejova E, Briscoe J. 2014. Integration of Signals along Orthogonal Axes of the Vertebrate Neural Tube Controls Progenitor Competence and Increases Cell Diversity ed. K.G. Storey. PLoS Biol 12: e1001907.

Spitz F, Furlong EEM. 2012. Transcription factors: from enhancer binding to developmental control. Nat Rev Genet 13: 613–626.

Thélie A, Desiderio S, Hanotel J, Quigley I, Van Driessche B, Rodari A, Borromeo MD, Kricha S, Lahaye F, Croce J, et al. 2015. *Prdm12* specifies V1 interneurons through cross-repressive interactions with *Dbx1* and *Nkx6* genes in *Xenopus*. Development 142: 3416–3428.

Timmer J, Chesnutt C, Niswander L. 2005. The Activin signaling pathway promotes differentiation of dI3 interneurons in the spinal neural tube. Developmental Biology 285: 1–10.

Tozer S, Le Dréau G, Marti E, Briscoe J. 2013. Temporal control of BMP signalling determines neuronal subtype identity in the dorsal neural tube. Development 140: 1467–1474.

Wagih O. 2017. ggseqlogo: a versatile R package for drawing sequence logos. Bioinformatics 33: 3645–3647.

Wang Y, Li X, Hu H. 2014. H3K4me2 reliably defines transcription factor binding regions in different cells. Genomics 103: 222–228.

Werner T, Hammer A, Wahlbuhl M, Bösl MR, Wegner M. 2007. Multiple conserved regulatory elements with overlapping functions determine So×10 expression in mouse embryogenesis. Nucleic Acids Res 35: 6526–6538.

Wickham H. 2016. Data Analysis. In ggplot2: Elegant Graphics for Data Analysis (ed. H. Wickham), pp. 189–201, Springer International Publishing, Cham 10.1007/978-3-319-24277-4_9 (Accessed January 20, 2025).

Zagorski M, Tabata Y, Brandenberg N, Lutolf MP, Tkačik G, Bollenbach T, Briscoe J, Kicheva A. 2017. Decoding of position in the developing neural tube from antiparallel morphogen gradients. Science 356: 1379–1383.

Zalc A, Rattenbach R, Auradé F, Cadot B, Relaix F. 2015. Pax3 and Pax7 Play Essential Safeguard Functions against Environmental Stress-Induced Birth Defects. Developmental Cell 33: 56–66.

Zechner D, Müller T, Wende H, Walther I, Taketo MM, Crenshaw EB, Treier M, Birchmeier W, Birchmeier C. 2007. Bmp and Wnt/β-catenin signals control expression of the transcription factor Olig3 and the specification of spinal cord neurons. Developmental Biology 303: 181–190.

Zhang Y, Liu T, Meyer CA, Eeckhoute J, Johnson DS, Bernstein BE, Nusbaum C, Myers RM, Brown M, Li W, et al. 2008. Model-based Analysis of ChIP-Seq (MACS). Genome Biology 9: R137.

Zhao Y, Vartak SV, Conte A, Wang X, Garcia DA, Stevens E, Kyoung Jung S, Kieffer-Kwon K-R, Vian L, Stodola T, et al. 2022. “Stripe” transcription factors provide accessibility to co-binding partners in mammalian genomes. Molecular Cell 82: 3398–3411.e11.

